# Bacterial protein function prediction via multimodal deep learning

**DOI:** 10.1101/2024.10.30.621035

**Authors:** Giulia Muzio, Michael Adamer, Leyden Fernandez, Lukas Miklautz, Karsten Borgwardt, Kemal Avican

## Abstract

Bacterial proteins are specialized with extensive functional diversity for survival in diverse and stressful environments. A significant portion of these proteins remains functionally uncharacterized, limiting our understanding of bacterial survival mechanisms. Hence, we developed Deep Expression STructure (DeepEST), a multimodal deep learning framework designed to accurately predict protein function in bacteria by assigning Gene Ontology (GO) terms. DeepEST comprises two modules: a multi-layer perceptron that takes gene expression and gene location as input features, and a protein structure-based predictor. Within DeepEST, we integrated these modules through a learnable weighted linear combination and introduced a novel masked loss function to fine-tune the structure-based predictor for bacterial species. These modeling choices are particularly well suited for bacteria due to the spatial organization of their circular genomes. Functionally related genes frequently co-localize and are co-transcribed within operons, allowing transcription dynamics to serve as crucial, condition-dependent regulatory signals. We show that DeepEST outperforms existing protein function prediction methods on a 25-species benchmark, relying solely on amino acid sequence or protein structure. Moreover, DeepEST predicts GO terms for unclassified hypothetical proteins across 25 human bacterial pathogens, facilitating the design of experimental setups for characterization studies. By combining expression, localization, and structure information in a unified deep learning framework, DeepEST bridges organism-specific data integration and structure-based transfer learning, providing a method tailored for bacterial protein function prediction in settings with structural and multi-condition expression data.

**Availability:** Accompanying code and data can be found under https://github.com/BorgwardtLab/DeepEST.

## 1 Introduction

A comprehensive characterization of protein function is a crucial aspect of understanding cellular biology. Despite this, a large majority of the known genes, even in extensively studied organisms, lack functional assessment for the encoded proteins [46]. This issue is even more pronounced in prokaryotes, where up to 60% of protein sequences remain unknown in terms of function [14]. Filling this knowledge gap is of crucial importance for advancements in various fields. For instance, it is essential for understanding bacterial physiology and their environmental adaptation. Moreover, it has the potential to drive advancements in biotechnological areas like bioremediation, bioenergy and novel medical treatments [3,13]. With roughly 10 million protein sequences of unknown function in bacteria [14], it is evident that experimental methods alone are insufficient to determine the function of each protein. Standard computational tools often rely on sequence similarity to annotate proteins, such as sequence alignment-based tools like BLAST [1,10], or on gene location-aware techniques [37]. More advanced methods utilize deep learning approaches, for example, DeepGOplus [27], which predicts protein functions by employing a convolutional neural network to automatically learn features from the amino acid sequences. However, despite their smaller and simpler genome, prokaryotes exhibit a high degree of functional redundancy and genetic diversity, making it challenging to predict protein functions based solely on sequences or gene location [4,37].

To overcome this challenge, it is necessary to integrate multiple data modalities that can capture vital yet different aspects of protein functionality, including protein structures, gene expression and gene location data. The information contained within protein structures provides valuable insights into their functional groups and domains [18,19]. DeepFRI [18], annotates proteins based on their structures. Besides protein structure, gene expression and location can be important descriptors, as genes involved in the same processes often exhibit similar expression patterns and they tend to be located in proximity within bacterial genomes [31]. While methods that combine different data sources already exist, they typically use protein sequences, protein structures, and protein-protein interaction data. Some examples include DeepGO [28], TransFun [7], HOPER [50] and the protein language model ProstT5 [20]. Furthermore, the majority of the methods in the literature have primarily been applied to eukaryote benchmark datasets and have rarely been used to study prokaryotes.

In this paper, we introduce DeepEST, a multimodal deep-learning approach specifically designed to predict protein functions in bacteria based on Gene Ontology (GO) annotations. DeepEST integrates structure, gene expression and location data as protein descriptors, offering the potential to enable accurate protein function prediction in bacteria. In bacteria, chromosomes are typically single and circular, so genes involved in related biological processes tend to be located close to each other on the chromosome and are often co-regulated. Moreover, many bacterial proteins are encoded in operons as polycistronic mRNAs, where multiple genes with related functions are transcribed together. DeepEST fine-tunes DeepFRI [18], by leveraging high-quality structural models available today [23,43,30] and incorporates an additional neural network using gene expression levels and locus features, capable of providing functional context. Throughout, we focus on bacterial genomes with (i) a high-quality reference genome/organization, (ii) protein structures (predicted or experimental), and (iii) multi-condition gene expression data. We evaluated DeepEST on a set of 25 bacterial species and predicted protein functions on a selection of ≈ 7,000 functionally unannotated hypothetical proteins.

## 2 Problem formulation

### 2.1 GO terms as label for protein function

To define the function of a protein, we utilize the GO terms [48], which are organized in three different ontologies, each represented as a directed acyclic graph (DAG). These ontologies include: Molecular Function, Cellular Components, and Biological Processes. Given the multifunctional nature of proteins and DAG structure of GO terms, a single protein can be associated with multiple GO terms simultaneously. Molecular Function terms are limited by precise semantics. We utilized Biological Process terms as a more comprehensive alternative to address this limitation. Biological Process encompasses a broader spectrum of protein activities, capturing diverse roles of proteins in cellular and physiological contexts. This holistic perspective accommodates proteins with multifaceted functions and integrates those whose specific molecular function remains elusive [41]. Given a set of *n*_p_ proteins 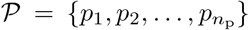 and a set of *n*_t_ terms 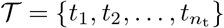, we define the matrix of the labels, 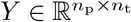, as

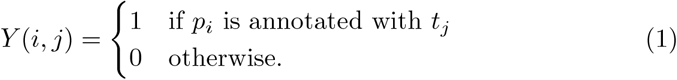

Hence, *Y* denotes the ground truth for protein function in the dataset we are studying, representing the GO term annotations within the set 𝒯, which consists of *n*_t_ unique GO terms annotated for at least one of the *n*_p_ proteins in 𝒫.

### 2.2 Prediction Problem

We use the combination of protein structures, gene expression and location data to enhance protein function prediction in bacteria. DeepEST predicts protein functions from a tuple of protein descriptors, *X* = (*X*_s_, *X*_e_). *X*_s_ represents the protein structures, while *X*_e_ comprises the gene expression and location data (Fig 1.a,b). Given the nature of the labels, we formulate the GO term prediction as a multi-modal and multi-label binary classification problem,

**Fig. 1:**
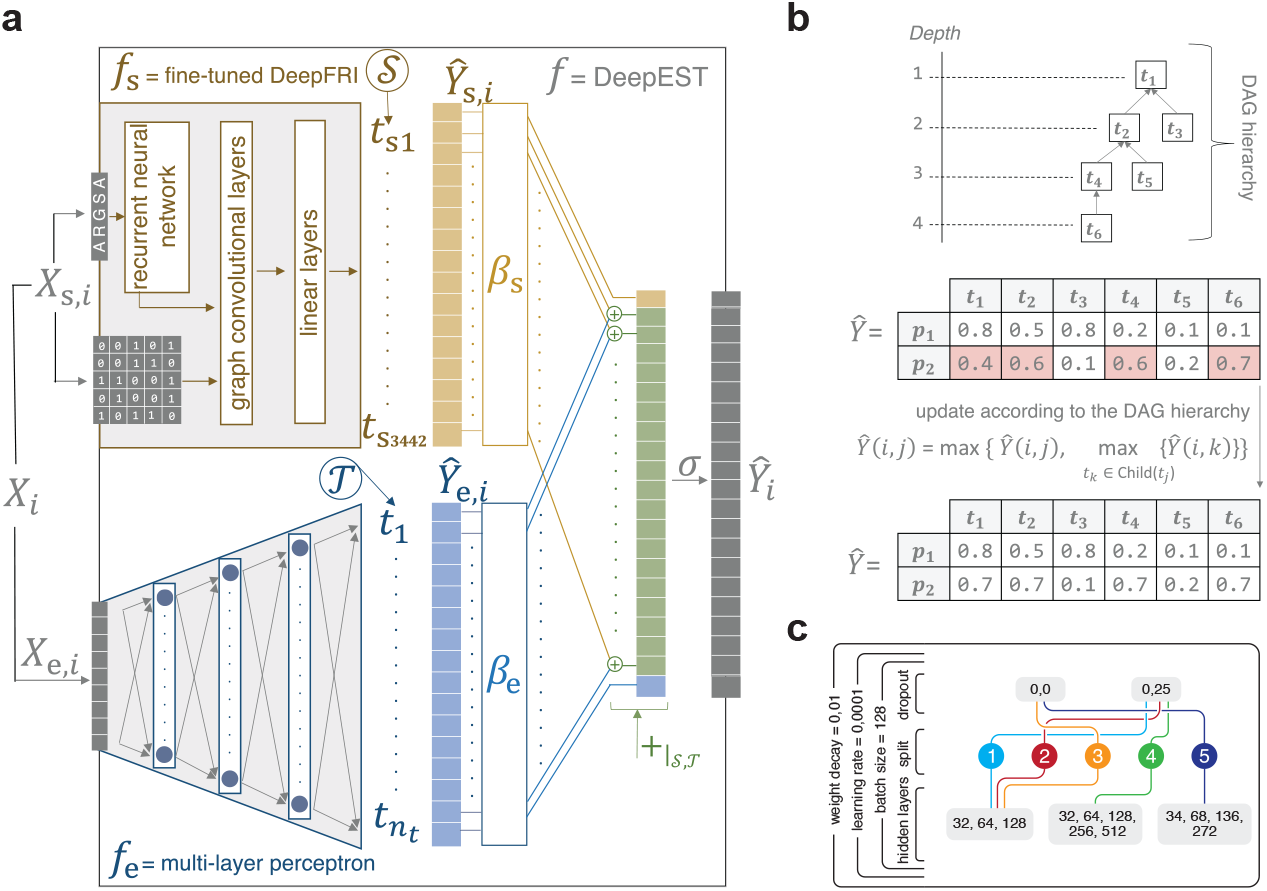
Schematic illustration of DeepEST framework. **a** Our approach is composed of two modules: (i) the expression-location module *f*_e_, a multi-layer perceptron, and (ii) the structure-based module *f*_s_, DeepFRI [18] exploited in a transfer learning setting. A protein *p*_*i*_ is represented by the tuple *X*_*i*_, composed of its expression-location features *X*_e,*i*_, and its protein structure *X*_s,*i*_, containing both the protein sequence and the adjacency matrix representing the amino acids contacts. *f*_e_ takes as input *X*_e,*i*_ and outputs *Ŷ*_e,*i*_ = *Ŷ*_e_(*i*, :), while *f*_s_ takes as input *X*_s,*i*_ and outputs *Ŷ*_s,*i*_ = *Ŷ*_s_(*i*, :). Predictions for protein *p*_*i*_, *Ŷ*_*i*_, are obtained by applying the sigmoid function (*σ*) on the masked linear combination (+_|*𝒮,𝒯*_ ) of *Ŷ*_s,*i*_ and *Ŷ*_e,*i*_. **b** At test step, predictions non-conforming the DAG hierarchy, visualized in red, are updated according to the rule from Eq. (9). **C** Hyperparameter configurations leading to the best validation performance for *f*_e_ in the five data splits.

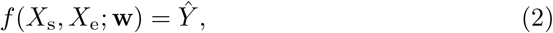

where the matrix of the predicted labels, 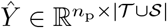, is obtained by applying *f* (·) to the input data, *X* = (*X*_s_, *X*_e_). *f* (·), the non-linear function parameterized in **w** representing DeepEST, is detailed in Methodology. Noteworthy, the matrix of the ground truth labels *Y* reports GO terms annotations for the terms in the set 𝒯, as introduced in the previous section. Instead, the matrix of the predicted labels *Ŷ* comprises predictions for the terms in |𝒯∪𝒮|, with 𝒮 being the set of terms predicted by the fine-tuned DeepFRI, i.e., the component of DeepEST that leverages protein structure data. The hyperparameter optimization for *f*_e_ is performed through grid search executed using W&B [6], with the best configurations shown in Fig 1.c. Additional details regarding hyperparameter optimization and training for each species are reported in S.1.

### 2.3 Metrics

To assess the quality of the predicted protein labels *Ŷ* relative to the ground truth annotations *Y*, we follow the evaluation protocol established by the Critical Assessment of Functional Annotation (CAFA) challenge [39]. In the term-centric setting, we compute the micro-averaged area under the precision–recall curve (*micro*-AUPRC), following [18]. For the protein-centric evaluation, we use the *F*_max_−score. Detailed definitions for all metrics are provided in S.2.

## 3 Methodology

In this section, we present DeepEST, describing its two distinct modules based on different data modalities and explaining their integration. We also outline how transfer learning is applied to enhance prediction accuracy.

### 3.1 Structure-based module

DeepFRI [18] predicts a set of 3, 442 GO terms in the biological process hierarchy, 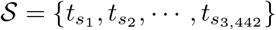, from the protein structures:

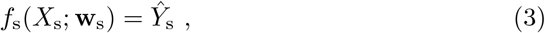

where *X*_s_ is constituted by the protein sequences and structures, *f*_s_(·) corresponds to DeepFRI and is parameterized in **w**_s_, and 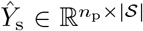 are the predicted labels. The architecture of *f*_s_(·) consists of two stages. Firstly, a recurrent neural network extracts representations from the protein sequences, generating a feature vector for each amino acid residue. Subsequently, a graph is constructed on an amino acid level, putting two residues into contact if they are less than 10Å apart. A protein is therefore represented through a graph with sequence-generated node features and is then fed through three graph convolutional network (GCN) layers [25]. Finally, the updated node features are concatenated, and the final predictions are computed with two linear layers. For the structure module of DeepEST, we harness DeepFRI in a transfer learning setting. Specifically, we keep the extracted features from the GCN layers and only retrain the weights of the final linear layers, henceforth denoted as **w**_s_. The structure of *f*_s_ is visualized at the top of Fig. 1.a. Contacts are derived from AlphaFold2 [43] (model version 4), and the resulting graph is passed to the GCN without pLDDT-based node/edge masking or edge reweighting.

### 3.2 Expression-location module

As motivated before, more accurate protein function prediction in bacteria requires a multimodal approach. Hence, our method adds another neural network, referred to as expression-location module, to the protein structure module. This module takes as input the experimentally measured gene expression values under different stress conditions and genetic locus descriptors (i.e., the gene’s *genomic locus* on the chromosome/plasmids, not protein subcellular localization), designed to capture the gene proximity information. Given that the bacterial genomes, with rare exceptions [21], consist of one main chromosome with a circular structure and smaller circular plasmids, we encode each position into polar coordinates. We then generate features by calculating the genetic distance, which involves calculating the sine and cosine of the angles formed by the positioning of the start and end positions of each gene. Lastly, we encode whether a gene is found on the main chromosome or on any of the plasmids, and the coding strand the gene is located on. The resulting gene expression *X*_expr_ and genome-location features *X*_loc_, with *n*_expr_ and *n*_loc_ number of features respectively, form a single tabular input for all *n*_p_ proteins, i.e., 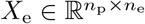 with

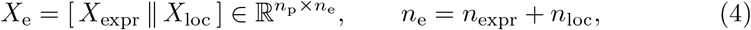

where *n*_e_ is the total number of features after concatenation and we denote horizontal concatenation with [ · ∥ ·]. Therefore, we use a multilayer perceptron (MLP) architecture to perform predictions according to this module,

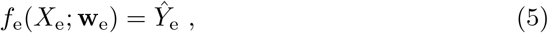

with *f*_e_(·) being the non-linear function representing the MLP and parameterized in **w**_e_, and 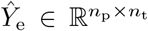 the matrix of predicted labels from the expression-location data. A visual representation of *f*_e_ is reported in the bottom of Fig. 1.a. Unlike the structure module, this module is trained *ab initio*.

### 3.3 Combining different modalities

DeepEST predicts protein functions by integrating the expression-location and structure data. This is achieved by performing a masked linear integration of Eq. (3) and Eq. (5), that allows to rewrite *f* (·), introduced in Eq. (2), as:

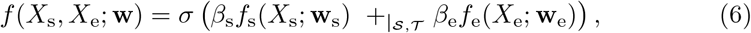

where *σ* is the sigmoid function. The learnable parameters of DeepEST denoted by **w**, include **w**_s_, **w**_e_, and the two integration parameters, *β*_s_ and *β*_e_. +_|*𝒮,𝒯*_ denote the masked sum performed accounting for the different sets of predicted GO terms. Indeed, exploiting *f*_s_(·) in a transfer learning setting means that the structure module predicts the set of terms originally predicted by DeepFRI, i.e. 𝒮. Instead, developing the expression-location module from scratch allows to predict the set 𝒯of GO terms annotations retrieved for the set of proteins 𝒫 in our dataset. Hence, when integrating the two modalities, DeepEST predicts terms in 𝒯 ∪ 𝒮, generating the output 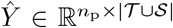. Explicitly, the masked linear integration of the two modalities writes *Ŷ* =

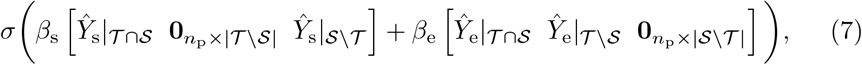

where *Ŷ*_e_|_(·)_ and *Ŷ*_s_|_(·)_ are the outputs of each module restricted to subsets of GO terms, namely the intersection of 𝒯 and 𝒮 (𝒯∩𝒮), the difference between 𝒯 and 𝒮 (𝒯 \ 𝒮), and the difference between 𝒮 and 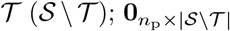 and 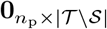 are the zero matrices with *n*_p_ rows and respectively |𝒮 \ 𝒯 | and |𝒯 \ 𝒮| columns; [· · ·] indicates the concatenation of three matrices. The output probabilities are given by sigmoid layer *σ*(·). This operation is visualized in Fig 1.a.

### 3.4 Transfer learning on the structure-based module

Transfer learning is a machine learning technique that harnesses the learned features from a pre-trained model to enhance the learning process for a closely related task, where usually only a dataset of small size is available. This is commonly accomplished by fine-tuning only the output layers of the model. We apply this technique on DeepFRI [18], to tailor the model to the bacterial organism we are investigating. To this end, we use a masked loss function *ℓ*(·),

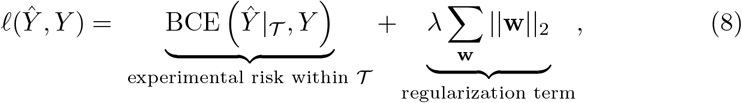

where *BCE*(·) is the multi-target binary cross-entropy loss, which calculates the experimental risk from the matrix of the predicted probabilities *Ŷ*, restricted to 𝒯, and the matrix of the ground truth labels *Y*; λ ∑_**w**_ ||**w**||_2_ is a weight decay regularization term with parameter λ [47]. The utilization of this loss allows for the transfer of label information for the set 𝒯 of GO terms in our dataset.

### 3.5 Update according to the DAG

Due to label dependence, any prediction needs to conform to the hierarchical structure of the GO terms. To this end, we use the common approach of updating the predictions at the test step according to the DAG [48], which follows the scheme

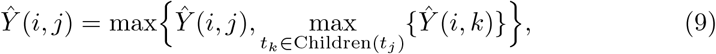

where Children(*t*_*j*_) represents the children of the term *t*_*j*_ according to the DAG. A visualization is reported in Fig. 1.b. Note that at training and validation time, we do not perform such updating steps to retain a differentiable loss function.

## 4 Experiments

We perform a comprehensive exploration of bacterial protein function prediction with a focus on 25 diverse human bacterial pathogens from different phylogenetic orders and Gram staining groups (Fig. S.1.a). We provide a detailed description of the data in S.3. In this section, we outline the experimental setup and present the results obtained from two analyses: (i) applying DeepEST to labeled data to assess performance, and (ii) predicting the functions of unlabeled proteins.

### 4.1 Evaluating the performance of DeepEST

#### Experimental setup

To assess the performance of DeepEST on predicting protein functions for the set of 25 bacterial species (Fig. S.1), we obtained the label matrix *Y* and the tuple of protein descriptors, *X* = (*X*_s_, *X*_e_), which includes protein structures and the expression-location features (preprocessing of expression features is described in S.4). We fine-tuned the pre-trained DeepFRI model, with the hyperparameter configuration identified in the original publication, as the structure module of DeepEST, *f*_s_. The MLP constituting the expression-location module, *f*_e_, is instead optimized and trained *ab initio*.

To minimize data leakage across train, validation, and test sets, we compared sequence- and structure-based splitting strategies (see S.5). Structure-based splitting yielded the least leakage and was therefore adopted as the default (see S.5.1). Accordingly, all main results in this paper use the Foldseek structure-based split, enforcing lDDT *<* 70% between training and validation/test proteins. To perform these operations and evaluate the performance of our model, we employ 5-fold nested cross-validation. For each pathogen, we assess the performance of DeepEST with structure- and sequence-based methods: DeepFRI[18], BLAST [1], DIAMOND [10], DeepGOCNN and DeepGOplus [27]. Further, we compare DeepEST to ProstT5[20], a state of the art transformer-based protein language model (pLM) that combines both structure and sequence information. All methods and the rationale behind their selection are described in S.6. Moreover, we assess the impact of DeepEST’s building blocks, the MLP constituting the expression-location module *f*_e_ and fine-tuned DeepFRI *f*_s_. We additionally evaluate DeepEST and DeepFRI with and without transfer learning.

#### DeepEST outperforms sequence and structure-based methods

To compare DeepEST with gene function annotation methods using different protein descriptors, we trained one model per species and compared to other methods. We used sequence-based traditional annotation methods (i.e., BLAST and Diamond), deep learning approaches (i.e., DeepGOCNN and DeepGOplus), a structure-based deep learning method, DeepFRI, and ProstT5, a pLM combining both sequence and structure. The *F*_max_ and *micro*-AUPRC scores described in Section 2.3 and S.2 were calculated for each species to assess performance.

DeepEST consistently outperforms all the sequence-only methods, i.e., BLAST, Diamond, DeepGOCNN, and DeepGOplus, both in terms of term-centric *micro-* AUPRC and protein-centric *F*_max_ Fig. 2 (a and b in Fig. S.9). This highlights the importance of considering more than just protein sequences when predicting protein function in bacteria. Instead, the use of DeepFRI, which utilizes protein structure data, already enables superior performance compared to the sequence-only baselines, showing the higher relative importance of protein structure over protein sequences for function prediction in bacteria. Moreover, DeepEST with usage of gene expression level and location data in addition to fine-tuning the structure-based module improved performance compared to DeepFRI, especially in terms of term-centric *micro-*AUPRC (Fig. 2 a and b). These results are confirmed by the additional metrics we utilized, namely term-centric *s*_min_, term-centric *macro*-AUPRC, and protein-centric AUPRC, which are introduced in S.2 and whose results are visualized in Fig S.10 and Fig S.11. These findings suggest that integrating the expression-location module along with the abundance of available protein structure data to refine the structure-based module allows for a more comprehensive analysis of protein functionality in bacteria.

**Fig. 2:**
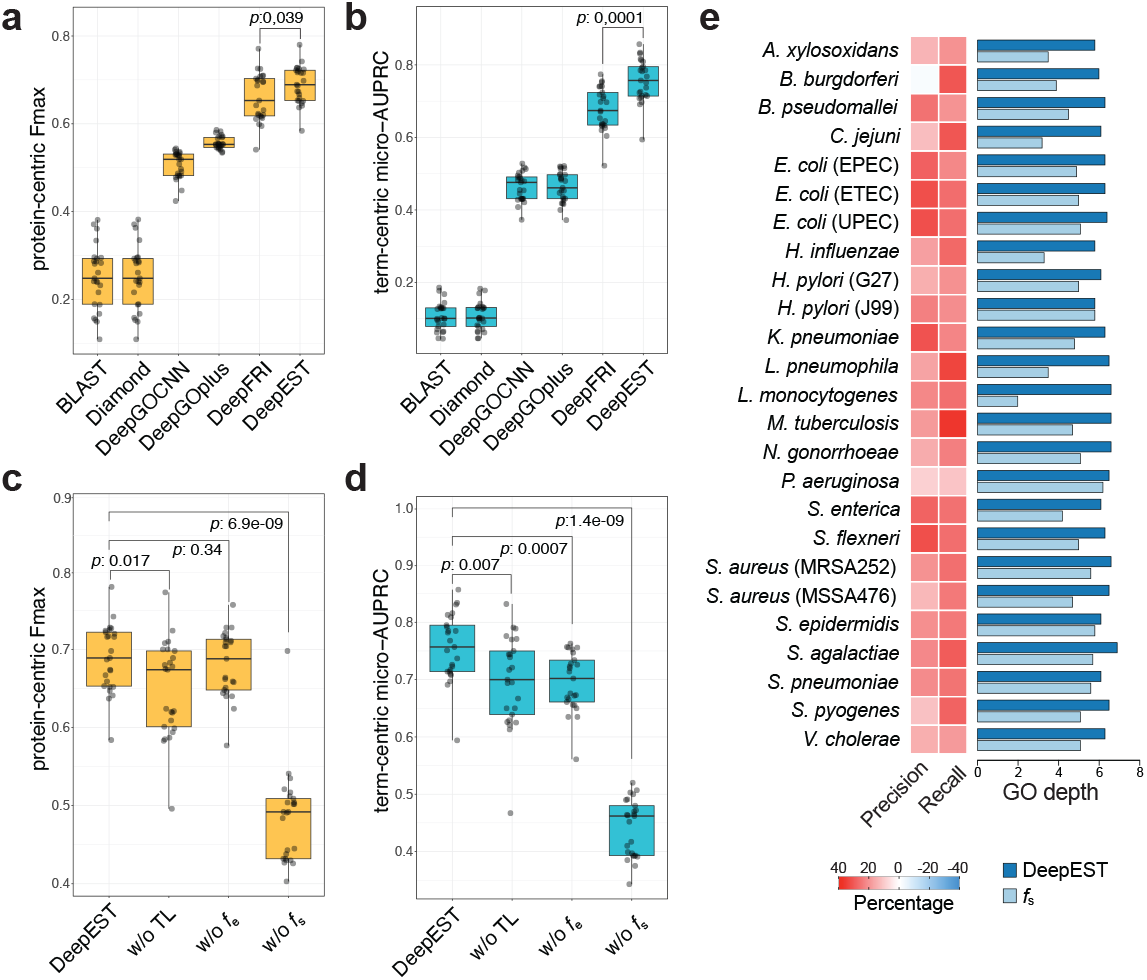
DeepEST outperforms established baselines in protein function predictions and achieves greater specificity in GO assignment by using expression location features. **a** The average protein-centric *F*_max_ and **b** term-centric *micro-*AUPRC values calculated for each species separately across the five sets for DeepEST and other methods: BLAST, Diamond, DeepGOCNN, DeepGOplus, and DeepFRI. **c** The average protein-centric *F*_max_ and **d** term-centric *micro-*AUPRC values calculated for each species separately across the five sets for DeepEST and DeepEST excluding one component at a time: (i) excluding our transfer learning (TL) technique, (ii) excluding the expression-location module, and (iii) excluding the structure module. **e** Comparison between DeepEST and *f*_s_ in terms of average %-change in precision and recall for each GO term across proteins (left panel), and average depth of the GO terms identified by either of the methods (right panel). Each dot in **a**,**b**,**c**,**d** represents *F*_max_ and *micro*-AUPRC scores for each species. *P-values* in **a** and **b** were calculated with Wilcoxon rank-sum test.

DeepEST consistently outperforms ProstT5 fine-tuned on the 25 bacterial species, both in terms of term-centric *micro-*AUPRC and protein-centric *F*_max_ Fig. 3. This highlights the importance of developing specialized models like DeepEST compared to using and fine-tuning more generalist models like ProstT5. We further demonstrate in Supplement S.8 that DeepEST empirically generalizes to a related CAFA5 strain, where PATHOgenex expression data is substituted with an independent stress-response expression compendium.

**Fig. 3:**
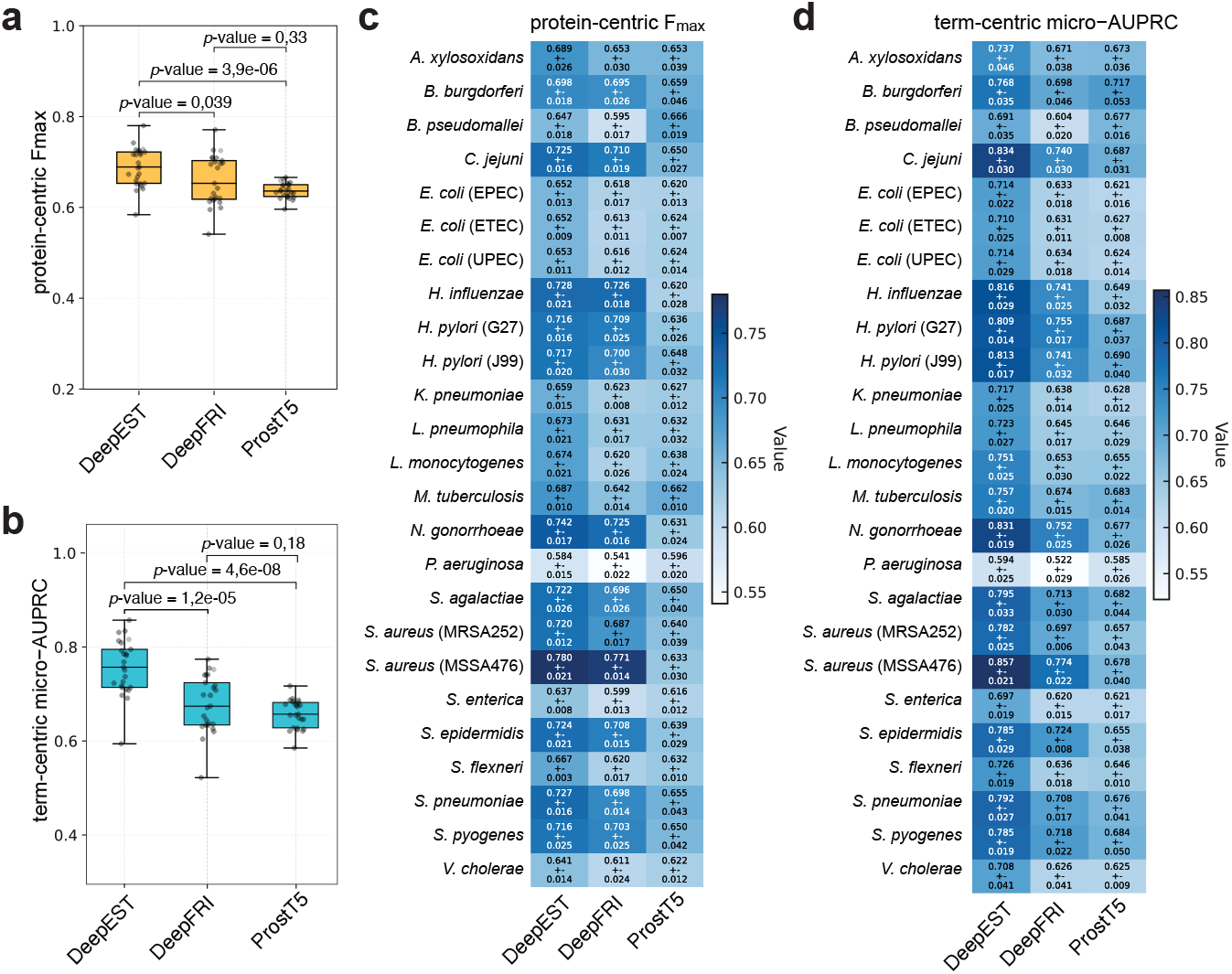
DeepEST outperforms protein language model ProstT5 in protein function predictions. **a** The average protein-centric *F*_max_ and **b** term-centric *micro-*AUPRC values calculated for each species separately across the five sets for DeepEST and other methods: DeepFRI and ProstT5. **c** Protein-centric *F*_max_ and **d** Term-centric *micro-*AUPRC values calculated for each species separately across the five sets for DeepEST, DeepFri and ProstT5 per each species. *P-values* in **a** and **b** were calculated with Wilcoxon rank-sum test.

#### Ablation study of DeepEST components

We assessed the contribution of DeepEST components by excluding transfer learning (TL), the structure-based module *f*_s_, or the expression-location module *f*_e_. TL consistently improved both protein-centric *F*_max_ and term-centric *micro-*AUPRC across species (Fig. 2c,d; Table S.1, Table S.2), and fine-tuning *f*_s_ further enhanced performance compared to DeepFRI (Fig. S.12). Removing *f*_e_ slightly affected *F*_max_ but reduced *micro-*AUPRC (Fig. 2c,d; Table S.2), indicating complementary information from expression-location features. DeepEST achieved higher average precision and recall for most species except *B. burgdorferi* and identified deeper GO terms (6.3 vs. 4.7 for *f*_s_; Fig. 2e; Table S.3). The unusual genomic organisation of *B. burgdorferi* with relatively small and linear chromosome and 21 plasmids (12 linear and 9 circular) underscore the impact of gene localization and expression on DeepEST’s prediction performance. The lower precision for *B. burgdorferi* than bacteria with simpler, predominantly circular chromosome and few plasmid genomes is not unexpected given 43 percent of genes located on 21 plasmids in *B. burgdorferi*. Excluding *f*_s_ led to a major decline in all metrics confirming structure data as the dominant contributor. Adding DeepGOCNN as additional protein descriptor provided no measurable gain (Fig. S.14).

### 4.2 Applying DeepEST on functionally unannotated proteins

Considering the diversity of protein function in bacteria and challenges in functional characterization of newly annotated proteins, a high percentage of bacterial proteins remains uncharacterized. These proteins are commonly known as hypothetical proteins, whose functions are unknown, and hence represent potential subjects for DeepEST. Within the 25 bacterial species, we identified 6,997 of hypothetical proteins that present both input features, *X*_s_ and *X*_e_, and remain functionally unannotated at the time of collecting GO terms annotations. We kept the optimal hyperparameter configurations listed in Fig 1.c for the expression-location module of DeepEST. The model, however, is retrained on the entire labeled dataset, setting aside a validation set for early stopping and to determine the optimal prediction threshold *τ*. Specifically, we choose the *τ* that maximizes the average F1-score calculated for each GO term across proteins, thereby optimizing the harmonic mean of precision and recall. Annotations are reported as the average of the five models identified in the five splits.

#### Results

DeepEST predictions yielded diverse tiers of the GO hierarchy for the 25 bacterial strains (available on our GitHub repository). At a GO depth greater than 5, DeepEST predicts 90 distinct GO terms, ranging from signal transduction and protein transport to aerobic respiration and RNA metabolism, for 727 gene products (Fig S.15). Most of these GO terms are associated with fewer than 25 gene products each, suggesting that our predictions are diverse and not biased towards fewer broader GO categories. We provide a detailed discussion on the biological relevance of these predictions in S.7.

## 5 Conclusion

We introduced **DeepEST**, a multimodal deep learning framework for bacterial protein function prediction. DeepEST integrates gene expression profiles and genomic location data, tailoring the model specifically for bacterial proteins, alongside protein structures to assign Gene Ontology (GO) terms. By combining organism-specific contextual data with structure-based transfer learning, DeepEST extends function prediction beyond sequence similarity approaches. As in most large-scale protein function prediction benchmarks, our GO/UniProt labels include computationally inferred annotations, introducing potential circularity where labels may derive from the input data. While this could inflate absolute performance metrics, relative comparisons across methods, as in our benchmarking experiments, should remain unaffected since all methods use identical labels.

With this caveat in mind, DeepEST outperforms sequence- and structure-based baselines, including DeepGOPlus [27], DeepFRI [18], BLAST [1], DIAMOND [10], and ProstT5 [20]. Across 25 bacterial pathogens, incorporating expression and location features yields more specific and deeper GO annotations than structure-only models. DeepEST further annotated nearly 7,000 previously uncharacterized proteins, revealing processes such as DNA repair and RNA metabolism and demonstrating its potential to guide targeted experimental validation. Overall, DeepEST provides a robust framework leveraging multimodal data and transfer learning to improve bacterial protein annotation and advance interpretable functional genomics in prokaryotes.

## Supplementary Material

### S.1 Hyperparameter optimization and training

The hyperparameter optimization for *f*_e_ is performed by means of grid search executed using W&B [6]. The hyperparameter optimized are listed in Table S.6. This hyperparameter search leads to having 192 potential combinations for each data split, 960 in total. The best configuration minimizes the multi-target binary cross-entropy loss on the validation set. Note that since we are using 5-fold nested cross-validation, we obtain a hyperparameter configuration per each of the five splits, as reported in Fig 1.c. Hence, this selection process is performed five times. However, to reduce computation and have a unified model per split across the 25 species, we perform the hyperparameter optimization on a unified dataset, which comprises all the proteins across the 25 species. Note that to avoid data leakage on this unified dataset as well, the structure-aware data splits generation using `foldseek` is performed on this dataset, to avoid very similar proteins from different species being in the training and test set at the same time, see details in S.5. Hence, the species-specific splits are subsequently obtained by sub-setting the unified dataset, as visualized in Figure 1. After having defined the best hyperparameter configurations across species, the sizes of the input and output are adapted to the number of expression-location features and GO terms accordingly. For details on the different sizes of the input and output, refer to the main text and Figure S.16 respectively.

DeepEST, which features the combination of the MLP with the best hyper-parameter configuration and DeepFRI subject to transfer learning, is trained on each species separately (ADAMW optimizer [32], cosine annealing with 10 epochs of linear warm up, base learning rate 0.0001, early-stopping based on validation loss with patience 4, maximum number of epochs 300). The reasoning behind the choice is that we did not observe any strong advantage in training across multiple species compared to training on each species separately.

### S.2 Evaluation metrics details

To assess the quality of the predicted protein labels *Ŷ* in comparison to the ground truth labels *Y*, we use the term-centric and protein-centric metrics introduced in Critical Assessment of Functional Annotation (CAFA) challenge [39]. These two categories of measures capture different aspects of the protein function prediction performance, namely the capability of the model to predict for each GO term what proteins are annotated with it, and its ability in predicting the function given a protein.

As for the term-centric metrics, we employ the *micro-* averaging area under the precision-recall curve (*micro*-AUPRC), following [18]. This metric considers all the GO terms simultaneously and is obtained by: (1) Flattening both *Y* and *Ŷ* to obtain two vectors of size *n*_p_*n*_t_, (2) Calculating the precision and recall of these vectors per each threshold *τ*, and (3) Obtaining the AUPRC given the precision and recall vectors. In regard to the protein-centric metric, we use the *F*_max_−score, defined as:

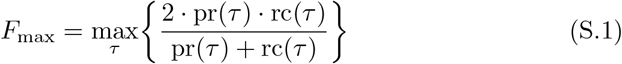

where *τ* ∈ [0, 1] represents some decision threshold; pr(*τ*) is obtained by averaging the precisions pr_*i*_(*τ*) across the proteins {*p*_*i*_ ∈ 𝒫 : *Ŷ* (*i, j*) ≥ *τ, j* = 1, · · ·, *n*_t_}; rc(*τ*) is the average of the recall rc_*i*_(*τ*) calculated per each protein *p*_*i*_ ∈ 𝒫. pr_*i*_(*τ*) and rc_*i*_(*τ*) are derived as follows:

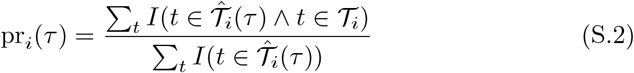

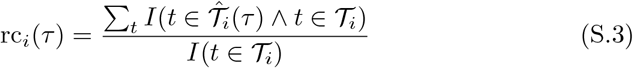

with 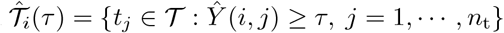 being the set of predicted GO terms for protein *p*_*i*_ at threshold *τ*, 𝒯_*i*_ = {*t*_*j*_ ∈ 𝒯 : *Y* (*i, j*) = 1, *j* = 1, · · ·, *n*_t_} the set of GO terms protein *p*_*i*_ is annotated with, *I*(·) the standard indicator function, and *t* a GO term.

In addition to *F*_max_ and *micro*-AUPRC, we complemented our analysis with additional metrics, namely term-centric minimum semantic distance (*s*_min_), term-centric macro-averaged area under the precision-recall curve (*macro*-AUPRC), and protein-centric area under the precision-recall curve (AUPRC).

The minimum semantic distance *s*_min_ is defined according to:

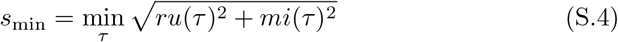

where *τ* ∈ [0, 1] is the decision threshold, *ru*(·) and *mi*(·) represent the uncertainty and the misinformation respectively and are calculated as follows:

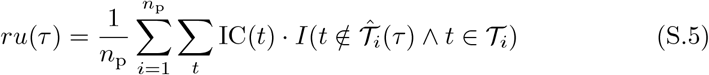

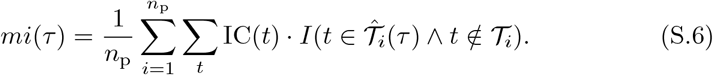

In the two equations above, *I*(·) is the standard indicator function, *t* a GO term, 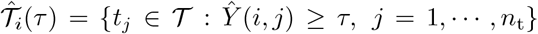 the set of predicted GO terms for protein *p*_*i*_ at threshold *τ*, 𝒯_*i*_ = {*t*_*j*_ ∈ 𝒯 : *Y* (*i, j*) = 1, *j* = 1, · · ·, *n*_t_} the set of GO terms protein *p*_*i*_ is annotated with, and IC(*t*) the information content of term *t*. The latter represents the term’s specificity by accounting for its frequency of annotations to the proteins in the dataset, and is calculated as *IC*(*t*) = −log_2_ *Pr*(*t*). *Pr*(*t*) is the probability of term *t* and is derived as the number of proteins annotated to *t* and its descendants, divided by the total number of annotated proteins.

The term-centric macro-averaged area under the precision-recall curve, *macro*-AUPRC, is derived by:

i. calculating precision and recall for each term *t* at for each threshold *τ* ∈ [0, 1];
ii. deriving the area under the precision-recall curve using the trapezoid rule for each term *t*;
iii. averaging these areas under the precision-recall curve across the terms.

The protein-centric area under the precision-recall curve, AUPRC, is derived by calculating pr(*τ*) and rc(*τ*) introduced in Eq. (S.2) for each threshold *τ*, and subsequently calculating the area under the pr(*τ*)-rc(*τ*) precision-recall curve by utilizing the trapezoid rule.

### S.3 Data

This section is devoted to introducing various data modalities utilized in our study, along with detailing the steps involved in data collection and preprocessing. Our analysis is specifically centered on the 25 bacterial species highlighted in Fig. S.1.a. For each of those, we conduct investigations using two datasets:

1. A labeled dataset, used to assess performance, for which we are able to collect the matrix of the ground truth labels, and
2. The dataset of hypothetical proteins, for which we are able to collect protein structures, gene expression and location features, while GO terms annotations are not available since these proteins have not been functionally characterized yet.

Concerning the labeled dataset, 32,573 proteins present both data modalities and GO term annotations. Instead, there are 6,997 hypothetical proteins for which both data views have been retrieved.

#### S.3.1 GO terms for protein function

As described in our results, we define protein function, i.e., the matrix of the ground truth labels *Y*, by utilizing GO terms annotations, with a specific focus on the Biological Process hierarchy. To collect available GO annotations, we query the UniProt database [11] (accessed on July 12, 2023) using the RefSeq protein identifier [36] of every known protein and the taxonomic reference code [15] of a given pathogen’s strain at hand. Note that the RefSeq protein identifier is a multi-species identifier and to obtain accurate annotations a taxonomy identifier has to be supplied. The number of unique GO terms obtained for each species can be found in Figure S.16. We used the GO ontology released on October 7, 2022 to retrieve a particular GO term’s children or ancestors. We do not filter GO terms by experimental evidence codes due to the lack of experimentally confirmed GO annotations, as shown in Figure S.17. The distribution of the number of annotated genes and the number of GO terms in each of the 25 species is visualized in Figure S.16. Note that we retain a GO term in case it is annotated for at least 10 proteins per species.

#### S.3.2 Protein structure data

The development of protein structure prediction methods, such as AlphaFold [23,43], and ESMFold [30], resulted in a wealth of high-quality protein structures across various species, including extensively investigated organisms as well as less studied species [43]. While experimentally confirmed protein structure may be available for some of the considered proteins, we only use structures in the AlphaFold database [43] in our approach, since obtaining all structures from one source mitigates the effect of covariate shifts in the input data. We query the AlphaFold database using the UniProt accession numbers of the proteins in question, thereby deriving the protein structure data *X*_s,*i*_ for all *p*_*i*_ ∈ 𝒫, i.e., for all the proteins we analyze, namely both the 32,573 proteins in the labeled dataset and the 6,997 hypothetical proteins.

#### S.3.3 Expression-location data

The expression-location features are derived from PATHOgenex dataset [4], which provides an in-depth global gene expression analysis across 11 host-mimicking stress conditions and control condition, listed in Fig. S.1.b. Specifically, as input features for our model, we consider the log-fold change values derived from the differential expression analysis of these 11 stress conditions compared to the control. As described before, we employed the engineered features derived from the genomic position and additional details on the preprocessing steps are reported Section S.4. As done for the protein structure data, expression-location features *X*_e,*i*_ are derived for all *p*_*i*_ ∈ 𝒫, namely for the 32,573 proteins in the labeled dataset and the 6,997 hypothetical proteins.

#### S.3.4 Visualization

The visualization of model performance was conducted using ggplot2 [44] in R version 4.2.2 [38]. Separate datasets were prepared for each evaluation metric, including micro-AUPRC and F-max. Each dataset was structured to categorize data by model type, enabling comparison across models. For each plot, jittered points were overlaid on box plots to visualize data dispersion. Statistical significance was assessed using the Wilcoxon rank-sum test to calculate p-values.

### S.4 Preprocessing of expression-location module

As described in the main paper, we utilize gene expression levels across various stress conditions (acidic stress, bile stress, low iron, nitrosative stress, oxidative stress, osmotic stress, nutritional downshift, hypoxia, stationary phase, temperature and virulence inducing condition, as shown in Fig. S.1.b) and location features as input to the expression-location module of DeepEST. As a preprocessing step for these features *X*_e_, we use `MinMaxScaler()` from `sklearn` to scale these features within the range [0, 1].

Note that *B. burgdorferi, B. pseudomallei, H. influenzae, H. pylori (G27), H. pylori (J99)*, and *M. tuberculosis* miss the measurements in one stress condition, while *L. pneumophila* lacks measurements in two stress conditions. When studying each species separately, these missing features are not considered. Instead, when analyzing multiple species simultaneously, these missing data are imputed with the average log fold expression under the same condition, across the other species.

### S.5 Importance of a proper evaluation framework

A commonly used strategy to generate data splits in protein function prediction foresees accounting for sequence similarity while generating the training/validation/test sets. Such strategy is aimed at diminishing possible data leakage from the training to the validation and test sets by clustering proteins with *high* sequence identity in the same fold. This allows to circumvent situations where proteins with highly similar amino acid sequences are present simultaneously in the training, validation and test set, thereby avoiding positively biased performance assessment. Such a strategy is necessary when calculating the performance of methods relying on amino acid sequences as input data, like many traditional tools such as BLAST [1].

Given the recent availability of protein structure data and methods exploiting such data, it becomes necessary to modify this paradigm to account for structure similarities as well, to avoid overly positive performance assessment. To estimate the effect of not accounting for protein structure similarity when evaluating the performance of a method that exploits structures, we apply DeepEST on splits generated while accounting for sequence similarity only. Highly similar proteins, that is, proteins presenting sequence identity higher or equal than 70%, are clustered together in either the training, validation, or test set. This is realized through the use of `CD-HIT` [17] clustering algorithm. Analogously to the pipeline applied to structure-based splits, this split generation procedure and the hyperparameter optimization are performed on the unified dataset generated by merging the data from the 25 species. The optimal configurations for the expression-location module are summarized in Table S.7 and Table S.8.

Results obtained according to the different splitting strategies are reported in Fig. S.4-S.5 and Fig.S.6-S.7, which present performance obtained with the structure-based and sequence-based data split procedure respectively. Note that since our study focuses on 25 different species, two different training strategies are possible, namely:

i. considering each species separately, thereby training 25 species-specific models, and
ii. considering all 25 species simultaneously, allowing to have a unique model trained on a unified dataset.

In this experiment, we consider and compare both scenarios. The performance of both DeepEST and fine-tuned DeepFRI is generally higher when training on data splits generated *not* accounting for structure similarity (i.e., *sequence*-based data splits) in comparison to accounting for it. This applies to both the species-specific and the unified analysis. A possible explanation lays in the fact that there is data leakage, i.e., very similar protein structures are present in the training, validation and test set simultaneously, therefore leading to overly positive performance assessment since the validation and the test set contain protein structures already seen during training. In addition, when structure similarity is accounted for when generating the splits, there is no evident benefit in training DeepEST and fined-tuned DeepFRI on multiple species simultaneously, as visualized in Fig. S.4 and Fig. S.5. In fact, *micro-*AUPRC and *F*_max_ result comparable (i.e., within one standard deviation from the average) between the two training strategies in 60% and 76% of the species, respectively. For DeepFRI, these percentages are 56% and 80% respectively. When structure similarity is not taken into account during the process of generating the splits, instead, it appears that training on all species simultaneously is beneficial for both DeepEST and fine-tuned DeepFRI. As reported in Fig. S.6 and Fig. S.7, there is an increase in performance in 97% of the cases. Given the previously reported observations, this increase in performance seems to be due to data leakage: very similar structures, from different species, appear in both training, validation and test set, since structure similarity is not considered when generating the data splits. Finally, the different data splits generation and training strategies have negligible impact on the expression-location module. In light of these observations, we retain as default splitting strategy the one accounting for structure similarity. With this choice, we can reduce the phenomenon of data leakage from the training to the validation and test set. Such considerations have been recently reported in [26]. Furthermore, we decide to train on each species separately, as there is no evident benefit in training on all species simultaneously. Note that these decisions are made on the results obtained on the validation sets.

#### S.5.1 Protein structure-based splitting improves generalizability of DeepEST

Given that DeepEST relies on a structure-based module, we generate the five data splits by accounting for protein structure similarity. This is accomplished by utilizing `foldseek` [24], which enables us to cluster proteins with high structure similarity (local distance difference test score [33] ≥ 70%) in the same fold (Fig S.2.a). This strategy is to ensure that the structures in the validation and test sets sufficiently differ from the ones in the training set to prevent any data leakage, enabling the evaluation of DeepEST’s generalizability to new structures [26], and thus, to previously unseen proteins. Therefore, we employed structure- and sequence-based splitting strategies to evaluate performance over data leakage, which is the case for the sequence-based splitting strategy. When splitting, we unified a dataset from all species and also generated data splits for single species by simply taking the subset belonging to that species from the unified training, validation and test sets respectively (Fig S.2.b). The performance of both DeepEST and DeepFRI in terms of *F*_max_ and *micro*-AUPRC is significantly higher when training on sequence-based splits, indicating a potential overperformance due to data leakage caused by similar protein structures in the split sets (Fig S.4, Fig S.5, Fig S.6, Fig S.7, Fig S.8). Noteworthy, this was more pronounced when using unified datasets, which are more likely to have similar protein structures. Hence, we compared the performance of the models with structure-based split datasets generated from unified data and subsets belonging to single species from the training, validation and test sets respectively. The analysis showed no significant difference in the performance of the models when using splits from all and single species datasets (Fig S.2.c-f and Fig S.8). This suggests improved generalizability of DeepEST by avoiding or minimizing similar structures in the training, validation, and test sets. We elaborate on the importance of employing this splitting strategy in the Supplement. Therefore, we applied structure-based splits.

The generation of the splits, which accounts for structure similarity between the proteins, is performed on the unified dataset. This is done to avoid data leakage from the training to the validation and test sets when performing hyperparameter optimization on the unified dataset, across the 25 species. In fact, performing the split generation on the single species and afterward unifying them would not assure that similar proteins from different species are not present in both training and test set of the same data split. Given the *i*-th data split of our five-fold nested cross validation obtained on the unified dataset, we generate the data splits for the single species by simply taking the subset belonging to that species from the training, validation and test sets respectively. Therefore, we employed structure- and sequence-based splitting strategies to evaluate performance over data leakage, which is the case for sequence-based splitting strategy.

### S.6 Selection and description of baselines

The additional baselines we use include methods taking protein sequences as input. We use traditional annotation methods (i.e., BLAST and Diamond) as well as deep learning-based approaches, such as ProstT5, DeepGOCNN and DeepGOplus. In the following, we provide a description of these tools.

- BLAST: the implementation of this baseline follows what done in [27]. Briefly, given a query protein from the validation or test set, its annotations are derived from the ones of the most similar protein in the training set, identified using sequence alignment algorithm BLAST [1] (we use its fastest implementation available under the Diamond tool [10]).
- Diamond: similarly to the baseline above, Diamond calculates annotation scores for a query protein by normalizing the sum of the alignment scores of similar proteins identified in the training set.
- DeepGOCNN: this is the convolutional part of DeepGOplus [27], and consists of several convolution filters, each with a different kernel size, to extract features directly from the input sequence. The resulting features are then pooled and concatenated, and given as input to a final linear layer with sigmoid activation that provides the output annotation scores.
- DeepGOplus: the annotations are obtained by weighting the predictions obtained using Diamond and DeepGOCNN.
- ProstT5: is a bilingual protein language model fine-tuned from ProtT5 to jointly learn amino acid sequences and structural representations encoded as 3Di tokens. Using a transformer encoder–decoder architecture, it translates bidirectionally between sequence and structure modalities, thereby capturing both evolutionary and geometric information. We employ ProstT5 by finetuning it on the GO terms. We fine-tuned a ProstT5 encoder with LoRA on the 25 bacterial species. Performance was assessed with GO term–centric *micro*-AUPRC and protein-centric *F*_max_ − score. Most importantly, we ran a learning-rate sweep at 10^−1^, 10^−2^, and 10^−3^, and 10^−2^. consistently gave the best results reported in the main manuscript (Fig 3).

Aside from the selected baselines, there is much recent work that has broadened protein function annotation beyond single-modality protein language models. Multimodal protein–text models aim to align protein representations with natural-language supervision and enable text-conditioned inference and retrieval (e.g., Evolla; [49]). In parallel, genome-scale language models trained on large genomic corpora (e.g., Evo2; [9]) provide long-context nucleotide representations that can support genome-informed functional inference. Complementing model-centric approaches, context-aware search/annotation platforms such as Gaia integrate genomic neighborhood information to improve microbial protein sequence search and annotation ([22]). As a PLM representative method, we used ProstT5 as a representative competitive baseline, which also performs strongly on public protein-function benchmarks like PROBE [42].

### S.7 Biological relevance of DeepEST predictions on functionally unannotated proteins

To further explore the biological significance of DeepEST-predicted GO terms with existing knowledge and literature, we conducted two case studies focusing on GO terms linked to DNA and RNA metabolism. For DNA metabolism, we analyzed “DNA repair” (GO:0006281) (Fig S.3a), while for RNA metabolism, we focused on “tRNA processing” (GO:0008033) and associated child terms (Fig S.3b). DeepEST identified genes potentially involved in DNA repair across various species with diverse taxonomic and physiological traits (Table S.4). DeepEST made high-confidence predictions for many genes containing Pfam domains such as AAA-ATPase, transcriptional regulator, and helix-turn-helix, all of which are associated with DNA binding and are typical of DNA repair proteins (Fig S.3c and Table S.4) [12], [2], [34]. Notably, DeepEST predicted GO terms for proteins with domains of unknown function also, indicating its prediction capability for proteins with unknown functional signatures (Fig S.3c Table S.4). In the case of tRNA processing, we identified domains linked to beta-lactamase superfamily proteins and sulfurtransferase TusA (Fig S.3d and Table S.5), which are critical for tRNA modification and maturation [5], [35], [40]. The term ‘tRNA processing’ was found across multiple species with notable GO term specificity in *Vibrio cholerae* (locus tag: VC2163). DeepEST predicted “tRNA wobble base modification” (GO:0002097) for *V. cholerae* protein containing TusA domain (Fig S.3 b and d, and Table S.5), consistent with previous literature [40]. In *Salmonella enterica* Typhimurium, we identified a gene with locus tag SL1344 RS05450, classified under “tRNA processing” (GO:0008033) and possesses domains linked to the beta-lactamase superfamily (Fig S.3d). Additionally, using the Co-PATHOgenex web app [16] and the same transcriptomic data as DeepEST, we identified a contiguous and co-expressed gene associated with the locus tag SL1344 RS05450, annotated as *iraM* Fig S.3e. IraM is an anti-adaptor protein that regulates the degradation of RpoS, a sigma factor essential for stress adaptation, and its expression is facilitated by rare tRNA production in *S. enterica* Typhimurium [8]. Through this reverse engineering exercise, we assessed the biological relevance of the predictions and the input data used by DeepEST. This empirical approach demonstrates that by integrating multiple layers of biological and biochemical data, DeepEST can improve both the accuracy of predictions and the precision of the information it provides.

### S.8 Using DeepEST with other transcriptomic datasets

DeepEST integrates structural features, genome location, and condition-specific gene expression to improve protein function prediction. To evaluate whether the model depends specifically on the PATHOgenex compendium used during training, we tested its performance using an independent transcriptomic dataset derived from a closely related strain and generated outside of the original database.

#### S.8.1 Cross-strain transfer to *Salmonella enterica* LT2

We selected *Salmonella enterica* serovar Typhimurium LT2, a strain included in the CAFA5 benchmark, and closely related to strain SL1344 used during training. Whole-genome alignment confirms near-complete collinearity and high nucleotide identity between LT2 and SL1344, with minimal rearrangements (Fig. S.18a). The dot-plot comparison (Fig. S.18a) shows a strong diagonal alignment across the entire chromosome, indicating conserved genome organization.

Because genome architecture is highly conserved between the two strains, positional genomic features learned from SL1344 can be reasonably transferred to LT2. In this experiment, we retained the genome-location encoding and substituted only the transcriptomic component.

#### S.8.2 Integration of an independent transcriptomic compendium

For LT2, we downloaded stress-response RNA-seq data from the Salmonella iModulon compendium. Expression values were processed to match the normalization strategy used for PATHOgenex data. Specifically, for each gene and condition, we computed Δ*z*-scores relative to exponential growth controls, thereby preserving the relative perturbation structure across conditions. Two configurations were evaluated:

1. 11 infection-relevant physiochemical conditions
2. 17 infection-relevant physiochemical conditions

These conditions were selected to reflect stress environments similar to those represented in PATHOgenex.

Genes present in SL1344 but absent in LT2 were imputed as knockout (expression value set to zero), ensuring consistent feature dimensionality. Importantly, the DeepEST model was not retrained. The pretrained model was directly applied with the substituted transcriptomic matrix.

#### S.8.3 Comparable performance across strains and datasets

Performance was evaluated using protein-centric F-max and GO term–centric micro-AUPRC across five cross-validation folds. As shown in Fig. S.18b, DeepEST achieves nearly identical performance when applied to LT2 with independently derived expression data compared to SL1344 using PATHOgenex.

- Protein-centric F-max remained approximately 0.65 across all configurations.
- GO term–centric micro-AUPRC remained approximately 0.71–0.72.
- Variability across folds (mean ± SD, n=5) overlapped between LT2 and SL1344.
- Notably, performance using 17 infection-relevant conditions was comparable to that obtained with 11 conditions, indicating that DeepEST does not require a large compendium of perturbations to remain effective.

#### S.8.4 Practical applicability and scope

These results demonstrate that DeepEST does not depend on a specific proprietary expression database. Instead, it can incorporate user-defined transcriptomic datasets provided that:

- A high-quality reference genome is available.
- Protein structures (predicted or experimental) can be obtained.
- Multi-condition gene expression measurements are available.
- Gene identifiers are mapped consistently.
- Genome organization is sufficiently conserved when transferring positional features across strains.

The LT2 experiment simulates a realistic application scenario in which a laboratory generates its own stress-response RNA-seq dataset and applies DeepEST without retraining the structural component. Under these conditions, performance remains stable and comparable to that obtained with the original training compendium. We emphasize that DeepEST is designed for bacteria where both structural and condition-specific expression data are available. It is not intended for metagenomic assemblies lacking genome organization or transcriptomic measurements. Within its intended scope, however, DeepEST provides a flexible multimodal framework capable of integrating independently generated transcriptomic datasets to enhance protein function prediction.

### S.9 Supplementary tables

**Table S.1:**
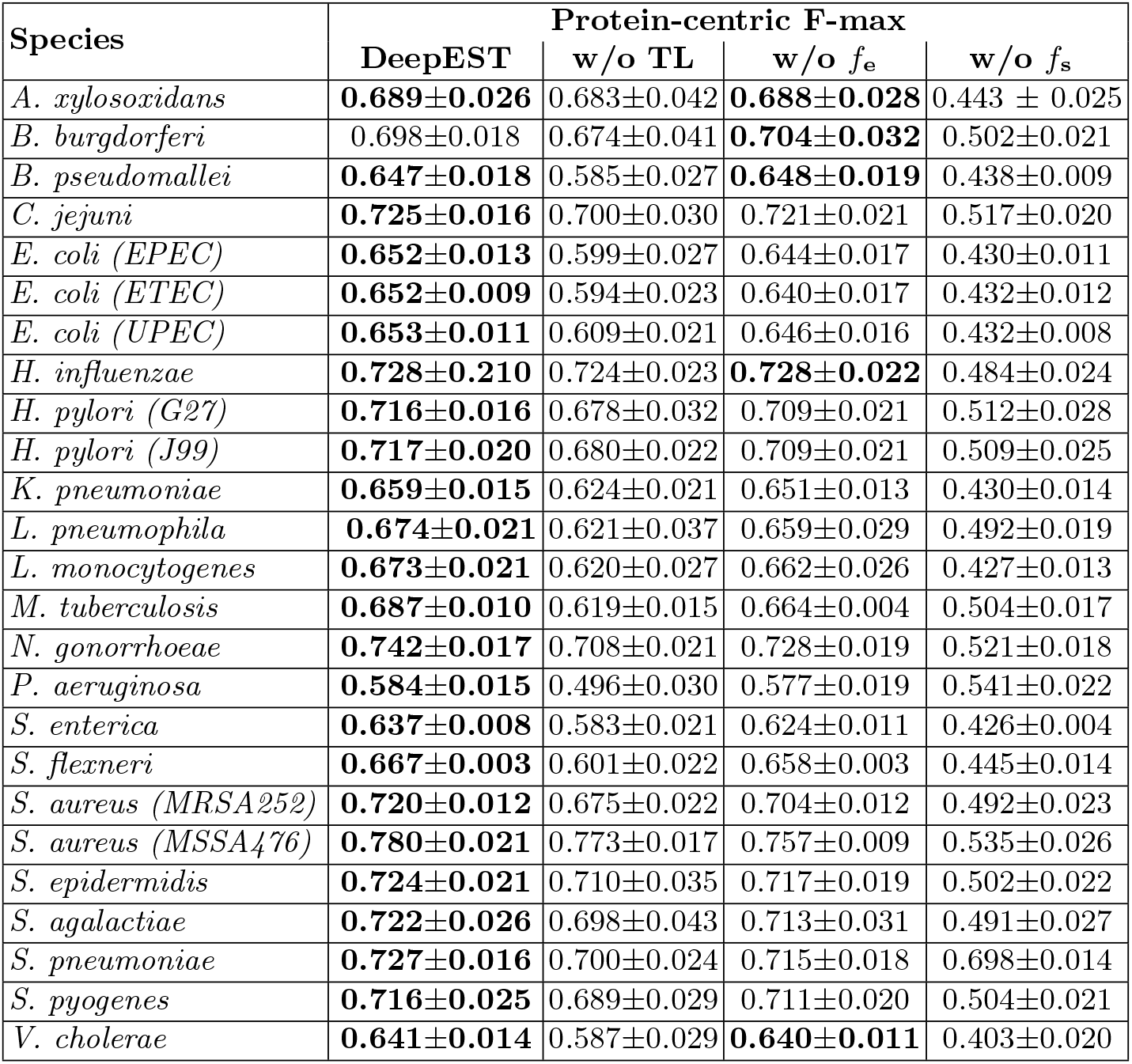
Assessment of the contributions of the components of DeepEST, by removing one at a time: (i) the use of our transfer learning (TL) technique, (ii) the expression-location module, and (iii) the structure module. The results are in terms of the average *F*_max_. ± standard deviation across the five test sets. In bold, we highlight the highest average score achieved per each species.

| Species | Protein-centric F-max |  |  |  |
| --- | --- | --- | --- | --- |
| | DeepEST | w/o TL | w/o $f_e$ | w/o $f_s$ |
| *A. xylosoxidans* | **0.689±0.026** | 0.683±0.042 | **0.688±0.028** | 0.443 ± 0.025 |
| *B. burgdorferi* | 0.698±0.018 | 0.674±0.041 | **0.704±0.032** | 0.502±0.021 |
| *B. pseudomallei* | **0.647±0.018** | 0.585±0.027 | **0.648±0.019** | 0.438±0.009 |
| *C. jejuni* | **0.725±0.016** | 0.700±0.030 | 0.721±0.021 | 0.517±0.020 |
| *E. coli (EPEC)* | **0.652±0.013** | 0.599±0.027 | 0.644±0.017 | 0.430±0.011 |
| *E. coli (ETEC)* | **0.652±0.009** | 0.594±0.023 | 0.640±0.017 | 0.432±0.012 |
| *E. coli (UPEC)* | **0.653±0.011** | 0.609±0.021 | 0.646±0.016 | 0.432±0.008 |
| *H. influenzae* | **0.728±0.210** | 0.724±0.023 | **0.728±0.022** | 0.484±0.024 |
| *H. pylori (G27)* | **0.716±0.016** | 0.678±0.032 | 0.709±0.021 | 0.512±0.028 |
| *H. pylori (J99)* | **0.717±0.020** | 0.680±0.022 | 0.709±0.021 | 0.509±0.025 |
| *K. pneumoniae* | **0.659±0.015** | 0.624±0.021 | 0.651±0.013 | 0.430±0.014 |
| *L. pneumophila* | **0.674±0.021** | 0.621±0.037 | 0.659±0.029 | 0.492±0.019 |
| *L. monocytogenes* | **0.673±0.021** | 0.620±0.027 | 0.662±0.026 | 0.427±0.013 |
| *M. tuberculosis* | **0.687±0.010** | 0.619±0.015 | 0.664±0.004 | 0.504±0.017 |
| *N. gonorrhoeae* | **0.742±0.017** | 0.708±0.021 | 0.728±0.019 | 0.521±0.018 |
| *P. aeruginosa* | **0.584±0.015** | 0.496±0.030 | 0.577±0.019 | 0.541±0.022 |
| *S. enterica* | **0.637±0.008** | 0.583±0.021 | 0.624±0.011 | 0.426±0.004 |
| *S. flexneri* | **0.667±0.003** | 0.601±0.022 | 0.658±0.003 | 0.445±0.014 |
| *S. aureus (MRSA252)* | **0.720±0.012** | 0.675±0.022 | 0.704±0.012 | 0.492±0.023 |
| *S. aureus (MSSA476)* | **0.780±0.021** | 0.773±0.017 | 0.757±0.009 | 0.535±0.026 |
| *S. epidermidis* | **0.724±0.021** | 0.710±0.035 | 0.717±0.019 | 0.502±0.022 |
| *S. agalactiae* | **0.722±0.026** | 0.698±0.043 | 0.713±0.031 | 0.491±0.027 |
| *S. pneumoniae* | **0.727±0.016** | 0.700±0.024 | 0.715±0.018 | 0.698±0.014 |
| *S. pyogenes* | **0.716±0.025** | 0.689±0.029 | 0.711±0.020 | 0.504±0.021 |
| *V. cholerae* | **0.641±0.014** | 0.587±0.029 | **0.640±0.011** | 0.403±0.020 |

**Table S.2:** Assessment of the contributions of the components of DeepEST, by removing one at a time: (i) the use of our transfer learning (TL) technique, (ii) the expression-location module, and (iii) the structure module. The results are in terms of the average *micro-*AUPRC ± standard deviation across the five test sets. In bold, we highlight the highest average score achieved per each species.

| Species | Term-centric AUPRC |  |  |  |
| --- | --- | --- | --- | --- |
| | DeepEST | w/o TL | w/o $f_e$ | w/o $f_s$ |
| *A. xylosoxidans* | **0.737±0.046** | 0.695±0.048 | 0.702±0.041 | 0.417±0.019 |
| *B. burgdorferi* | **0.768±0.035** | 0.721±0.035 | 0.723±0.049 | 0.500±0.029 |
| *B. pseudomallei* | **0.691±0.035** | 0.626±0.024 | 0.635±0.030 | 0.397±0.012 |
| *C. jejuni* | **0.834±0.030** | 0.789±0.020 | 0.756±0.035 | 0.491±0.027 |
| *E. coli (EPEC)* | **0.714±0.022** | 0.621±0.023 | 0.666±0.023 | 0.391±0.015 |
| *E. coli (ETEC)* | **0.710±0.025** | 0.625±0.021 | 0.656±0.025 | 0.393±0.017 |
| *E. coli (UPEC)* | **0.714±0.029** | 0.639±0.019 | 0.661±0.026 | 0.392±0.015 |
| *H. influenzae* | **0.816±0.029** | 0.776±0.030 | 0.757±0.034 | 0.465±0.025 |
| *H. pylori (G27)* | **0.809±0.014** | 0.771±0.014 | 0.747±0.046 | 0.503±0.036 |
| *H. pylori (J99)* | **0.813±0.017** | 0.770±0.018 | 0.753±0.039 | 0.490±0.033 |
| *K. pneumoniae* | **0.717±0.025** | 0.650±0.020 | 0.655±0.021 | 0.393±0.017 |
| *L. pneumophila* | **0.751±0.025** | 0.700±0.025 | 0.676±0.034 | 0.452±0.019 |
| *L. monocytogenes* | **0.723±0.027** | 0.667±0.023 | 0.669±0.035 | 0.399±0.014 |
| *M. tuberculosis* | **0.757±0.020** | 0.694±0.011 | 0.673±0.009 | 0.470±0.017 |
| *N. gonorrhoeae* | **0.831±0.019** | 0.791±0.017 | 0.763±0.018 | 0.507±0.023 |
| *P. aeruginosa* | **0.594±0.025** | 0.467±0.027 | 0.561±0.027 | 0.343±0.010 |
| *S. enterica* | **0.697±0.019** | 0.629±0.021 | 0.635±0.021 | 0.385±0.008 |
| *S. flexneri* | **0.726±0.019** | 0.650±0.017 | 0.672±0.008 | 0.410±0.013 |
| *S. aureus (MRSA252)* | **0.782±0.025** | 0.717±0.027 | 0.714±0.031 | 0.463±0.026 |
| *S. aureus (MSSA476)* | **0.857±0.021** | 0.832±0.017 | 0.756±0.038 | 0.520±0.034 |
| *S. epidermidis* | **0.785±0.029** | 0.744±0.021 | 0.728±0.029 | 0.472±0.026 |
| *S. agalactiae* | **0.795±0.033** | 0.750±0.038 | 0.734±0.054 | 0.462±0.035 |
| *S. pneumoniae* | **0.792±0.027** | 0.744±0.027 | 0.726±0.028 | 0.464±0.019 |
| *S. pyogenes* | **0.785±0.019** | 0.739±0.027 | 0.720±0.031 | 0.480±0.035 |
| *V. cholerae* | **0.708±0.041** | 0.613±0.042 | 0.650±0.041 | 0.375±0.019 |

**Table S.3:** Comparison between *f*_s_ (i.e., fine-tuned DeepFRI) and DeepEST in terms of percentage change in precision, recall and F1-score calculated for each GO term across proteins. These annotations are generated by thresholding the annotation scores using the threshold that maximizes the F1-score on the validation set. We furthermore report the average and median of the GO terms that only DeepEST is capable to correctly retrieving, shown under “Depth DeepEST”, in comparison to the average and median GO terms that both methods are capable to identify, under “Depth intersection”, and to the average and median depth of the GO terms that solely *f*_s_ was capable of correctly identifying, i.e. “Depth *f*_s_”.

| Species | % change<br>vs. $f_s$ | | | Depth<br>DeepEST | | Depth<br>intersec. | | Depth<br>$f_s$ | |
| --- | --- | --- | --- | --- | --- | --- | --- | --- | --- |
|  | Prec. | Recall | F1 | Av. | Med. | Av. | Med. | Av. | Med. |
| *A. xylosoxidans* | +14.1 | +20.5 | +19.9 | 5.8 | 5.0 | 4.7 | 4.0 | 3.5 | 4.0 |
| *B. burgdorferi* | -1.9 | +30.8 | +16.3 | 6.0 | 5.0 | 4.2 | 4.0 | 3.9 | 4.0 |
| *B. pseudomallei* | +25.8 | +20.2 | +27.4 | 6.3 | 6.0 | 5.1 | 5.0 | 4.5 | 4.5 |
| *C. jejuni* | +12.5 | +30.8 | +24.7 | 6.1 | 5.0 | 4.8 | 4.0 | 3.2 | 3.5 |
| *E. coli (EPEC)* | +28.8 | +22.1 | +29.0 | 6.3 | 6.0 | 5.0 | 5.0 | 4.9 | 5.0 |
| *E. coli (ETEC)* | +32.1 | +26.7 | +33.5 | 6.3 | 6.0 | 5.0 | 5.0 | 5.0 | 5.0 |
| *E. coli (UPEC)* | +32.0 | +26.5 | +33.0 | 6.4 | 6.0 | 5.0 | 5.0 | 5.1 | 5.0 |
| *H. influenzae* | +17.9 | +27.5 | +26.4 | 5.8 | 6.0 | 4.8 | 4.0 | 3.3 | 3.0 |
| *H. pylori (G27)* | +15.2 | +20.7 | +22.1 | 6.1 | 6.0 | 4.5 | 4.0 | 5.0 | 5.0 |
| *H. pylori (J99)* | +23.5 | +21.2 | +26.3 | 5.8 | 5.5 | 4.6 | 4.0 | 5.8 | 7.0 |
| *K. pneumoniae* | +31.4 | +22.7 | +31.5 | 6.3 | 6.0 | 5.0 | 5.0 | 4.8 | 5.0 |
| *L. pneumophila* | +17.4 | +34.0 | +27.4 | 6.5 | 6.0 | 5.0 | 5.0 | 3.5 | 3.5 |
| *L. monocytogenes* | +22.2 | +26.4 | +27.9 | 6.6 | 7.0 | 5.2 | 5.0 | 2.0 | 2.0 |
| *M. tuberculosis* | +18.9 | +36.7 | +32.4 | 6.6 | 7.0 | 4.9 | 4.0 | 4.7 | 4.0 |
| *N. gonorrhoeae* | +15.8 | +23.7 | +24.5 | 6.6 | 7.0 | 4.8 | 4.0 | 5.1 | 4.0 |
| *P. aeruginosa* | +8.9 | +11.1 | +12.0 | 6.5 | 6.0 | 5.4 | 5.0 | 6.2 | 6.0 |
| *S. enterica* | +28.4 | +26.0 | +32.2 | 6.1 | 6.0 | 5.1 | 5.0 | 4.2 | 4.0 |
| *S. flexneri* | +32.1 | +26.5 | +34.1 | 6.3 | 6.0 | 5.0 | 5.0 | 5.0 | 5.0 |
| *S. aureus (MRSA252)* | +20.5 | +26.3 | +27.2 | 6.6 | 7.0 | 5.0 | 5.0 | 5.6 | 7.0 |
| *S. aureus (MSSA476)* | +13.5 | +24.6 | +24.0 | 6.5 | 7.0 | 4.9 | 4.0 | 4.7 | 4.5 |
| *S. epidermidis* | +21.0 | +24.6 | +27.1 | 6.1 | 6.0 | 4.9 | 4.0 | 5.8 | 5.5 |
| *S. agalactiae* | +22.0 | +29.8 | +29.8 | 6.9 | 8.0 | 4.8 | 4.0 | 5.7 | 5.0 |
| *S. pneumoniae* | +21.6 | +25.6 | +27.5 | 6.1 | 6.0 | 4.9 | 4.0 | 5.6 | 5.0 |
| *S. pyogenes* | +11.9 | +28.4 | +22.2 | 6.5 | 7.0 | 4.7 | 4.0 | 5.1 | 5.0 |
| *V. cholerae* | +15.2 | +18.8 | +20.8 | 6.3 | 6.0 | 5.2 | 5.0 | 5.1 | 5.0 |

**Table S.4:**
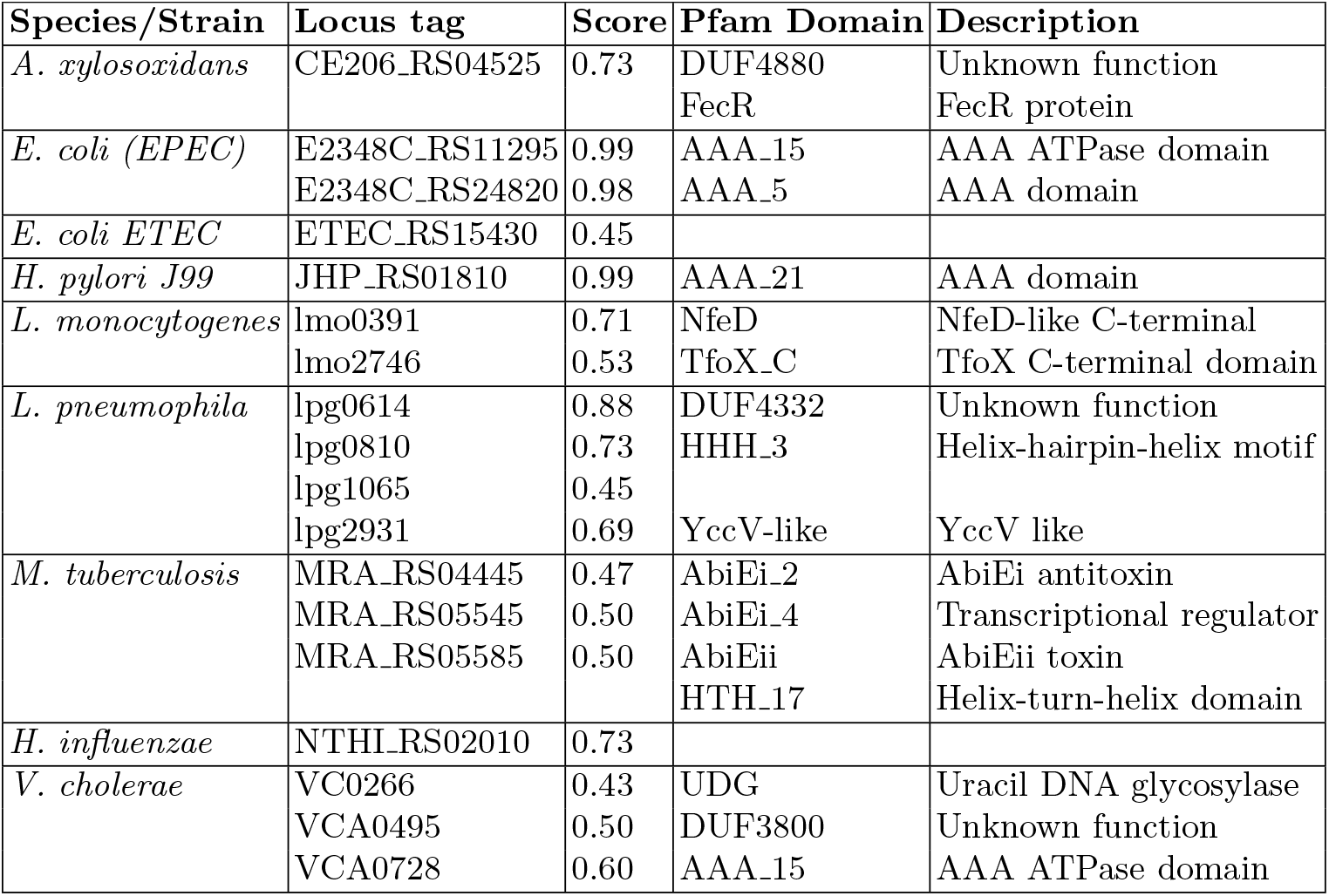
Genes with GO:0006281 (DNA repair)

**Table S.5:** Genes with GO:0008033 (tRNA processing)

| Species/Strain | Locus tag | Score | Pfam Domain | Description |
| --- | --- | --- | --- | --- |
| *B. burgdorferi* | BB_O17 | 0.74 | DUF693 | Unknown function |
| *M. tuberculosis* | MRA_RS04100 | 0.50 | DUF429 | Unknown function |
| *S. enterica* | SL1344_RS05450 | 0.37 | Lactamase_B_2 | Beta-lactamase |
|  | SL1344_RS14220 | 0.42 | DUF4435 | Unknown function |
| *V. cholerae* | VC0605 | 0.69 | UPF0231 | Uncharacterised protein |
|  | VC2163 | 0.84 | TusA | Sulfurtransferase TusA |
|  | VCA0475 | 0.34 | Fic | Fic/DOC family |

**Table S.6:** To choose the best hyperparameter configuration for the experimental module (i.e., the MLP), we perform a grid search using W&B. The hyperparameters and the values we tried are reported in this table, leading to 192 different combinations for each data split (960 total runs). We perform this hyperparameter optimization on both the structure-based and sequence-based splits.

| Hyperparameter | values |
| --- | --- |
| hidden layers | - [32, 64, 128, 256, 512]<br>- [34, 68, 136, 272]<br>- [32, 64, 128]<br>- [32, 64]<br>- [512, 16]<br>- [512, 16, 512]<br>- [32, 256, 64, 512]<br>- [32, 128, 32, 128] |
| dropout | - 0.0<br>- 0.25 |
| activation | ReLU |
| batch size | - 32<br>- 64<br>- 128 |
| base learning rate | - 0.01<br>- 0.0001 |
| weight decay | - 0.0<br>- 0.01 |

**Table S.7:** Best hyperparameter configurations identified for the MLP in the five *sequence*-based data splits.

| Split | hidden layers | dropout | batch size | learning rate | weight decay |
| --- | --- | --- | --- | --- | --- |
| 1 | [32, 128, 32, 128] | 0.25 | 32 | 0.0001 | 0.01 |
| 2 | [32, 128, 32, 128] | 0.25 | 32 | 0.0001 | 0.01 |
| 3 | [32, 128, 32, 128] | 0.25 | 32 | 0.0001 | 0.01 |
| 4 | [32, 64, 128, 256, 512] | 0.25 | 32 | 0.0001 | 0.01 |
| 5 | [32, 128, 32, 128] | 0.25 | 32 | 0.0001 | 0.01 |

**Table S.8:** Sizes of the input and output layers for the MLP for each species.

| Species | Input size | Output size |
| --- | --- | --- |
| all species | 17 | 1711 |
| *A. xylosoxidans* | 17 | 239 |
| *B. burgdorferi* | 16 | 159 |
| *B. pseudomallei* | 16 | 387 |
| *C. jejuni* | 17 | 210 |
| *E. coli (EPEC)* | 17 | 445 |
| *E. coli (ETEC)* | 17 | 435 |
| *E. coli (UPEC)* | 17 | 416 |
| *H. influenzae* | 16 | 242 |
| *H. pylori (G27)* | 16 | 179 |
| *H. pylori (J99)* | 16 | 203 |
| *K. pneumoniae* | 17 | 404 |
| *L. pneumophila* | 15 | 289 |
| *L. monocytogenes* | 17 | 376 |
| *M. tuberculosis* | 16 | 301 |
| *N. gonorrhoeae* | 17 | 233 |
| *P. aeruginosa* | 17 | 694 |
| *S. enterica* | 17 | 449 |
| *S. flexneri* | 17 | 420 |
| *S. aureus (MRSA252)* | 17 | 291 |
| *S. aureus (MSSA476)* | 17 | 198 |
| *S. epidermidis* | 17 | 271 |
| *S. agalactiae* | 17 | 229 |
| *S. pneumoniae* | 17 | 238 |
| *S. pyogenes* | 17 | 204 |
| *V. cholerae* | 17 | 474 |

### S.10 Supplementary figures

**Fig. S.1:**
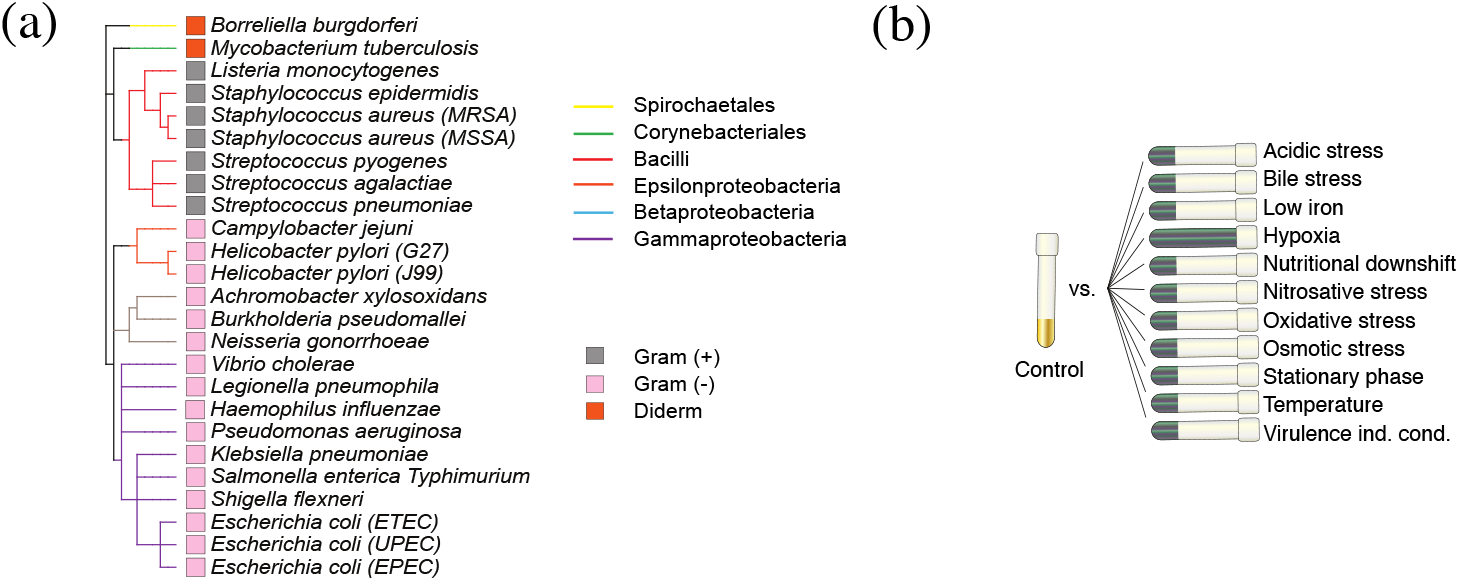
Overview of the bacteria and expression features utilized in this study: Phylogenetic clustering of the set of 25 selected strains (obtained using PhyloT in Newick [45] format and represented using iTol [29]), and (b) the experimental setup of differential expression dataset from Avican et al. [4].

**Fig. S.2:**
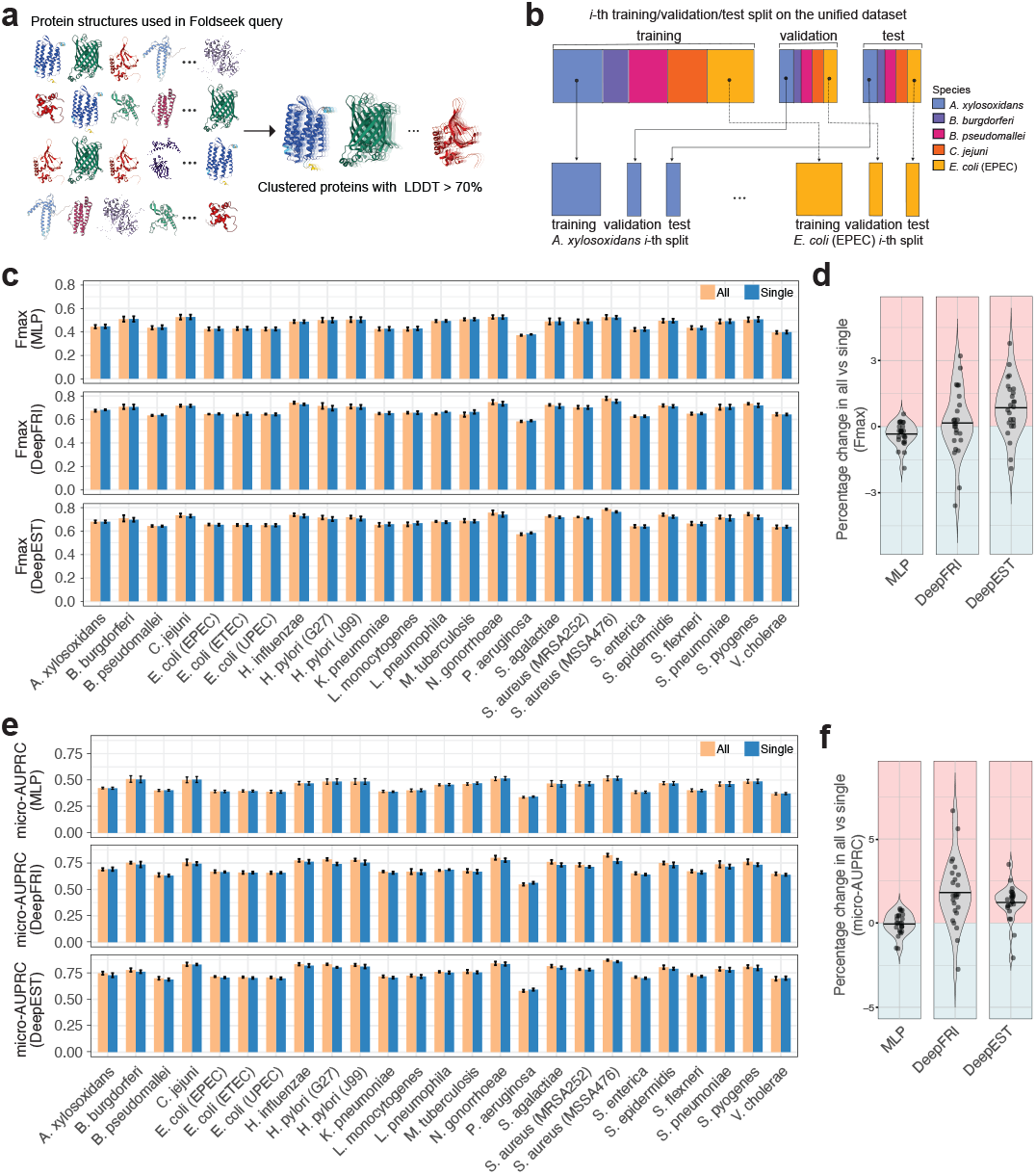
Accounting structure similarity with a unified dataset from 25 species or a single species exhibits similar performance suggesting no data leakage in training, validation, and test sets. **a** Available protein structures from the 25 bacterial species retrieved from AlphaFold were used as queries in Foldseek to cluster proteins with high structure similarity local distance difference test score (LDDT) ≥ 70%.**b** Visual representation of split generation for a representative subset of five species. The generation of the splits, which account for structure similarity between the proteins, is performed on the unified dataset. **c** Comparison of training MLP (i.e., *f*_e_), fine-tuned DeepFRI (i.e., *f*_s_) and DeepEST on each species separately (“single”) and on the unified dataset comprised of the 25 species (“all”) in terms of mean *F*_max_ on the five *validation* sets, with *structure*-based splits. Each panel corresponds to a specific model used in the analysis. The bar plot shows the mean *F*_max_ values for protein function prediction in each species. The colors represent datasets, namely “single” and “all”, used in training. **d** Percentage change of mean *F*_max_ values when training the models according to the “all” strategy in contrast to the “single”. **e** Comparison of training the three models in terms of average *micro*-AUPRC with same strategy explained in **c. f** Percentage change of mean *micro*-AUPRC values when training the models according to the “all” strategy in contrast to the “single”. The error bars in **c** and **e** represent the standard deviation for each species. Each dot in **d** and **f** represents a percentage change of *F*_max_ and *micro*-AUPRC, respectively, for each species and strait lines represent the mean values.

**Fig. S.3:**
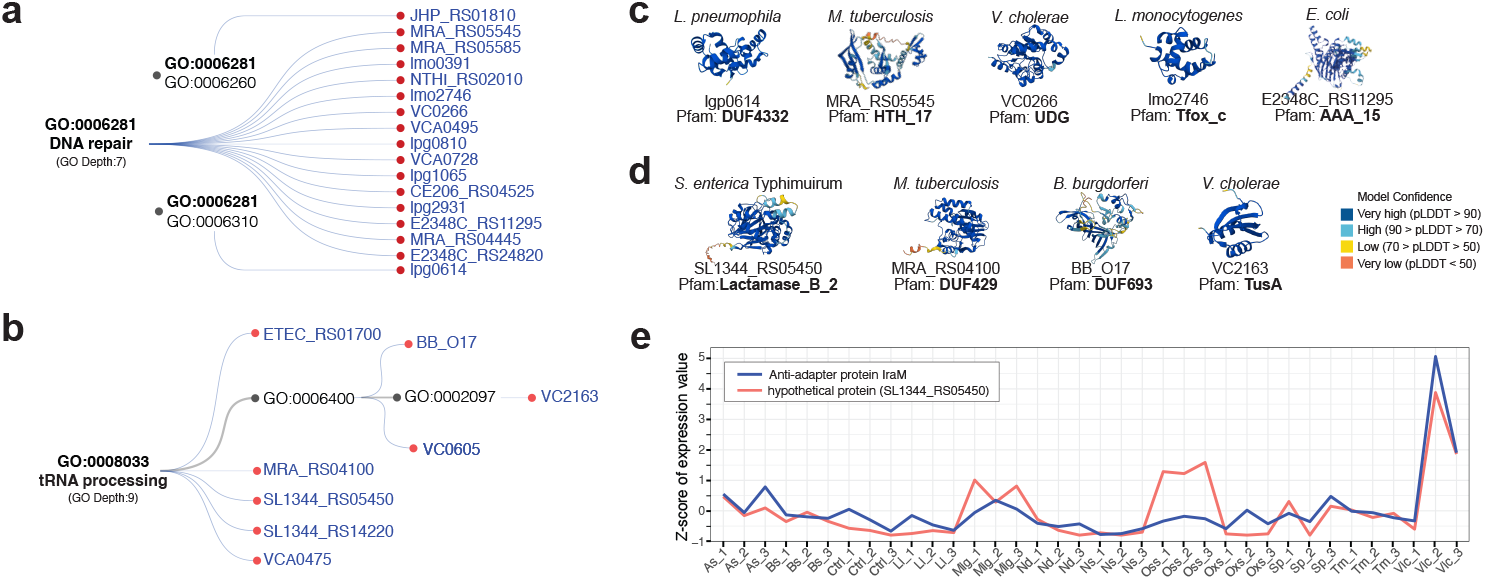
DeepEST predicts functions via assigning specific biological process GO terms to bacterial hypothetical proteins with domains of known and unknown functions. Ontology tree of DeepEST-assigned **a** DNA repair (GO:00066281) with GO hierarchy depth of 7 and **b** tRNA processing (GO:0008033) with its child nodes, along with its predicted proteins from different bacterial species. Representatives of proteins with domains of known and unknown functions together with Alphafold-predicted structures for proteins **c** assigned with DNA repair and **d** tRNA processing. **e** Expression patterns of the gene encoding anti-adaptor protein IraM and SL1344 RS05450 encoding a hypothetical protein in *S. enterica Typhimurium* under each replicate of the 12 conditions. Abbreviations of the conditions in **e** are as follows: As; Acidic stress, Bs; Bile stress, Ctrl; Control, Li; Low iron, Mig; Microaerophilic growth, Nd; Nutritional downshift, Oss; Osmotic stress, Oxs; Oxidative stress, Sp; Stationary phase, Tm; Temperature; Vic. Virulence inducing condition.

**Fig. S.4:**
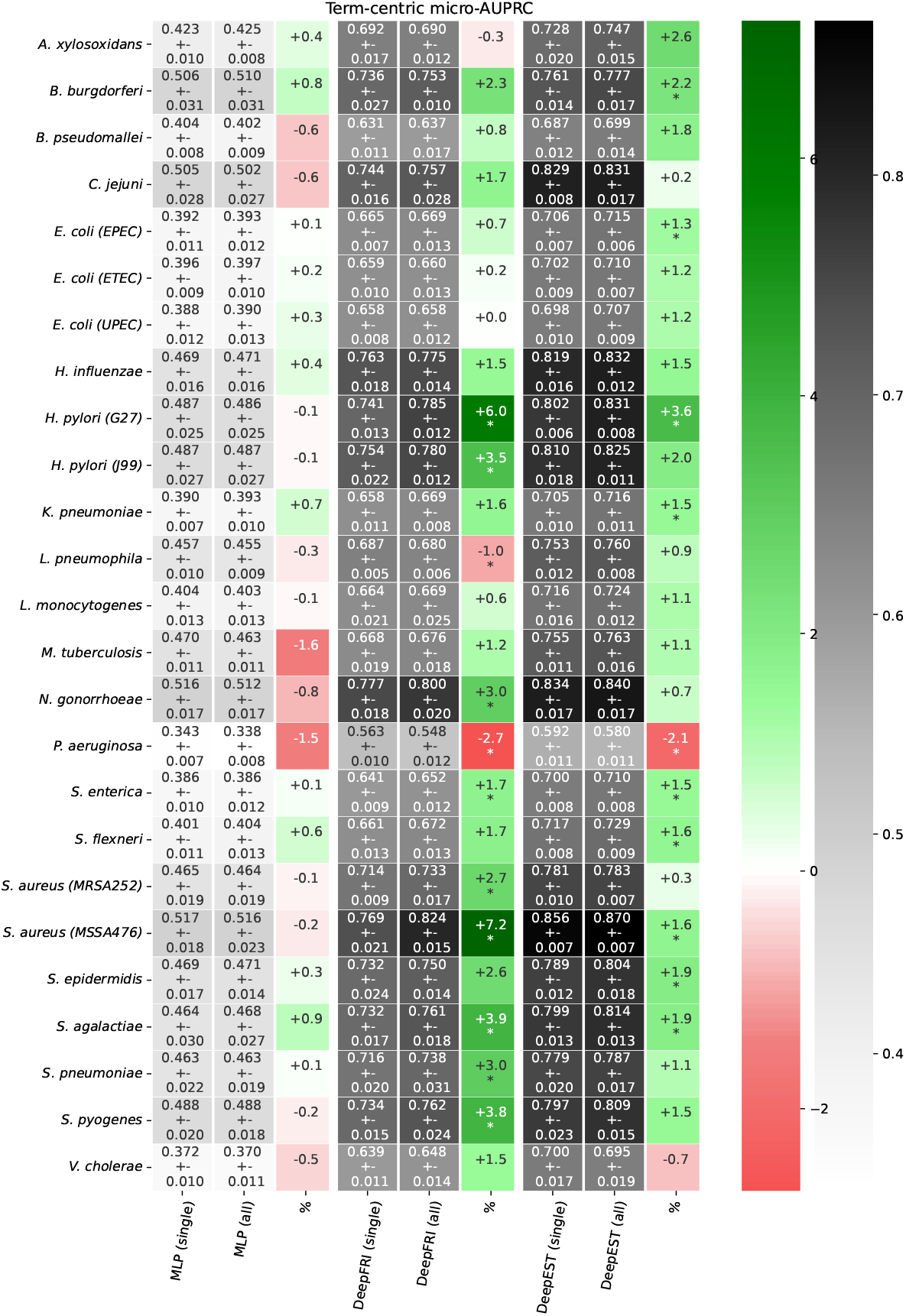
Comparison between training MLP (i.e., *f*_e_), fine-tuned DeepFRI (i.e., *f*_s_) and DeepEST on each species separately (“single”) and on the unified dataset comprised of the 25 species (“all”) in terms of average *micro*-AUPRC on the five *validation* sets, with *structure*-based splits. Each model presents three columns, namely “single”, “all” and %. The latter is the percentage change obtained when training according to the “single” strategy in contrast to the “all”, with an “*” when the difference between the two averages exceeds one standard deviation. Each column represents a species.

**Fig. S.5:**
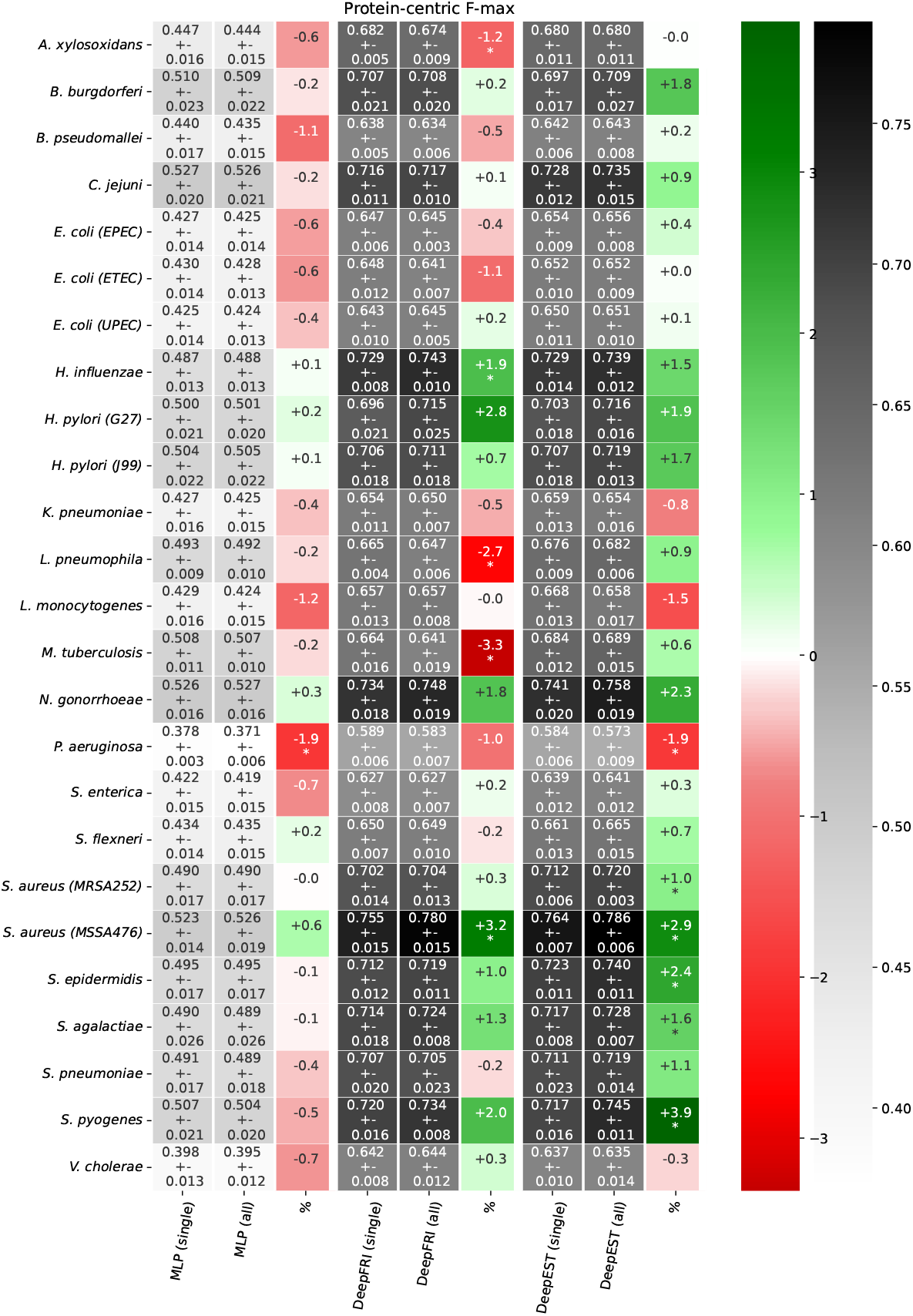
Comparison between training MLP (i.e., *f*_e_), fine-tuned DeepFRI (i.e., *f*_s_) and DeepEST on each species separately (“single”) and on the unified dataset comprised of the 25 species (“all”) in terms of average *F*_max_ on the five *validation* sets, with *structure*-based splits. Each model presents three columns, namely “single”, “all” and %. The latter is the percentage change obtained when training according to the “single” strategy in contrast to the “all”, with an “*” when the difference between the two averages exceeds one standard deviation. Each column represents a species.

**Fig. S.6:**
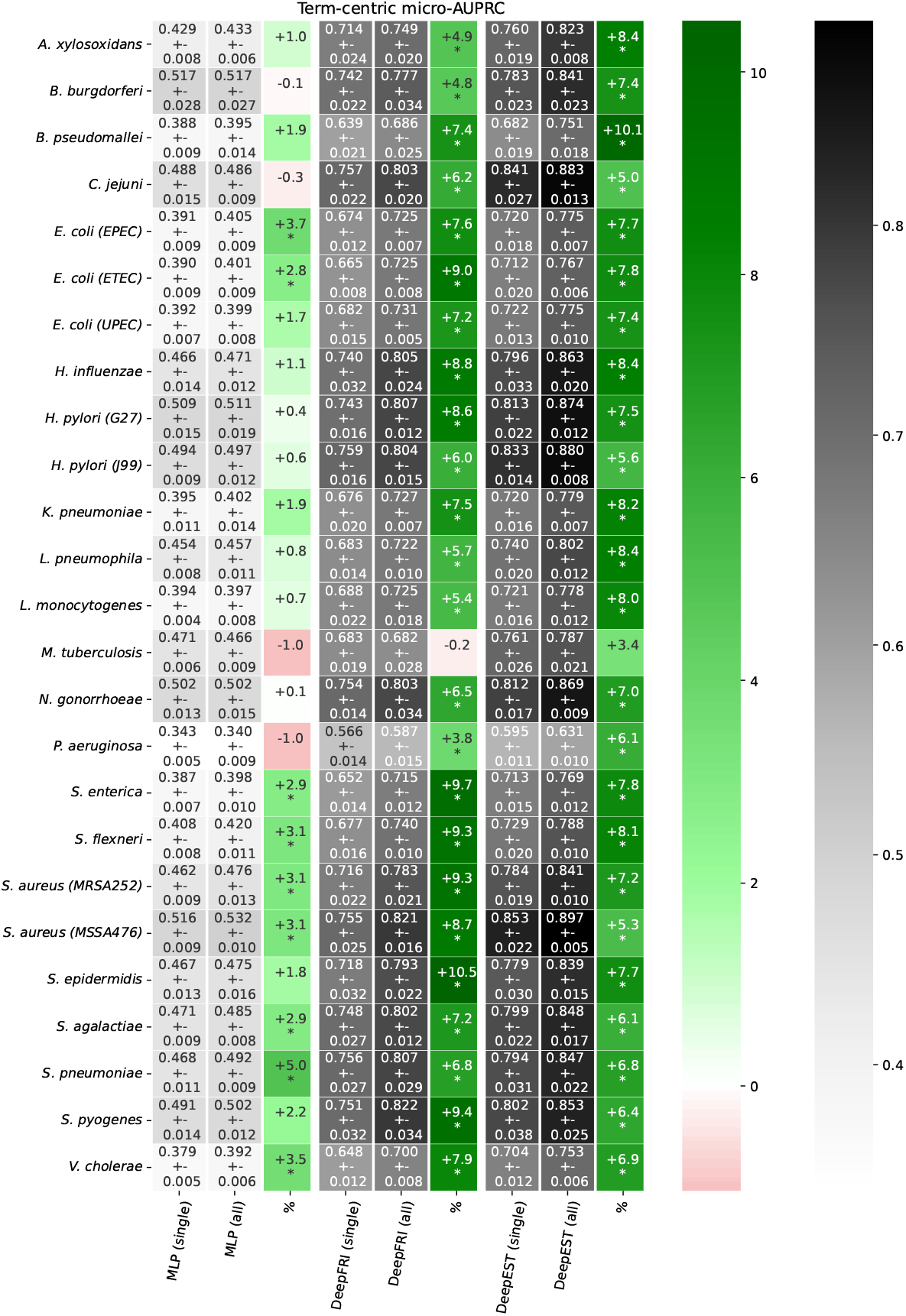
Comparison between training MLP (i.e., *f*_e_), fine-tuned DeepFRI (i.e., *f*_s_) and DeepEST on each species separately (“single”) and on the unified dataset comprised of the 25 species (“all”) in terms of average *micro*-AUPRC on the five *validation* sets, with *sequence*-based splits. Each model presents three columns, namely “single”, “all” and %. The latter is the percentage change obtained when training according to the “single” strategy in contrast to the “all”, with an “*” when the difference between the two averages exceeds one standard deviation. Each column represents a species.

**Fig. S.7:**
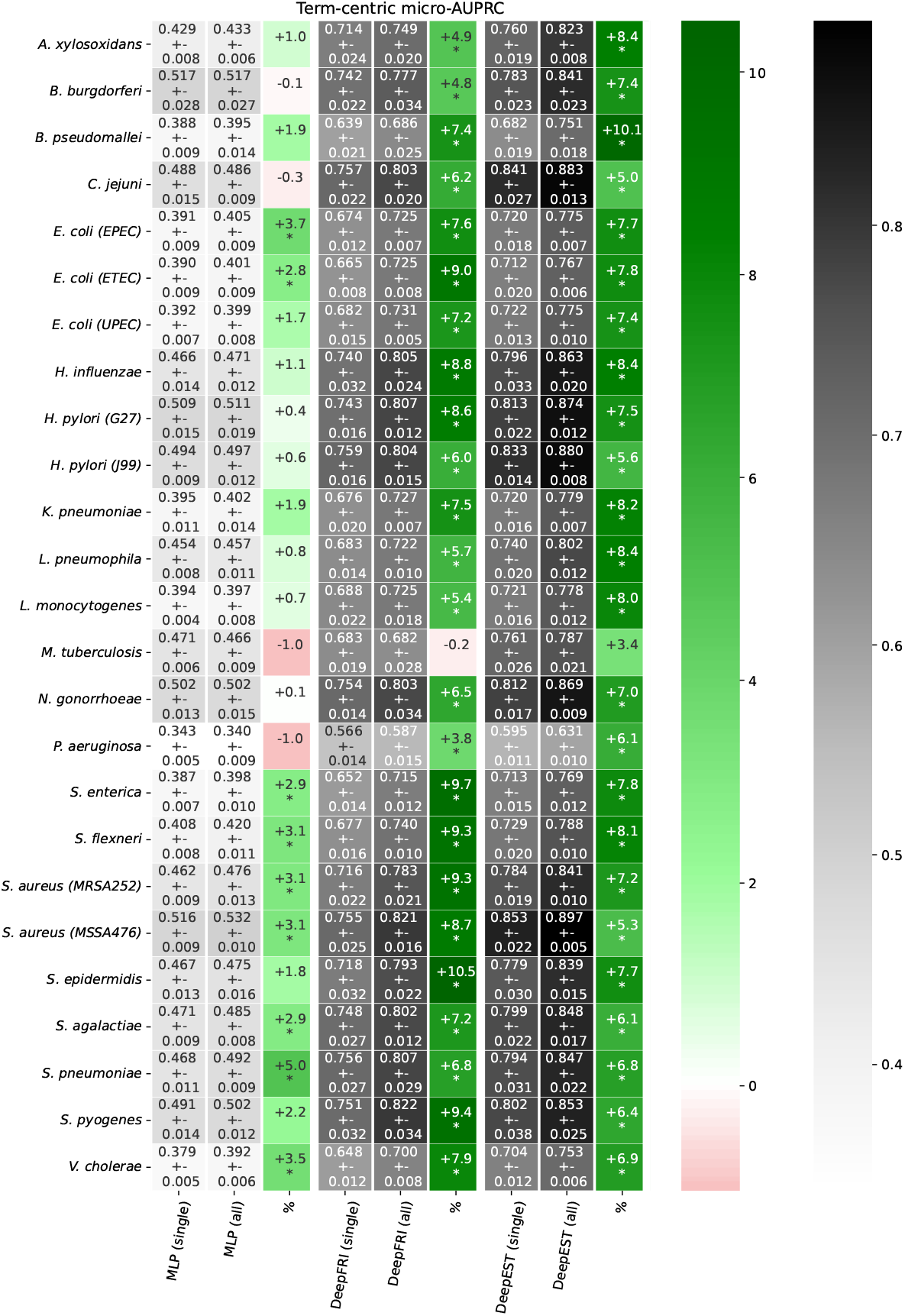
Comparison between training MLP (i.e., *f*_e_), fine-tuned DeepFRI (i.e., *f*_s_) and DeepEST on each species separately (“single”) and on the unified dataset comprised of the 25 species (“all”) in terms of average *F*_max_ on the five *validation* sets, with *sequence*-based splits. Each model presents three columns, namely “single”, “all” and %. The latter is the percentage change obtained when training according to the “single” strategy in contrast to the “all”, with an “*” when the difference between the two averages exceeds one standard deviation. Each column represents a species.

**Fig. S.8:**
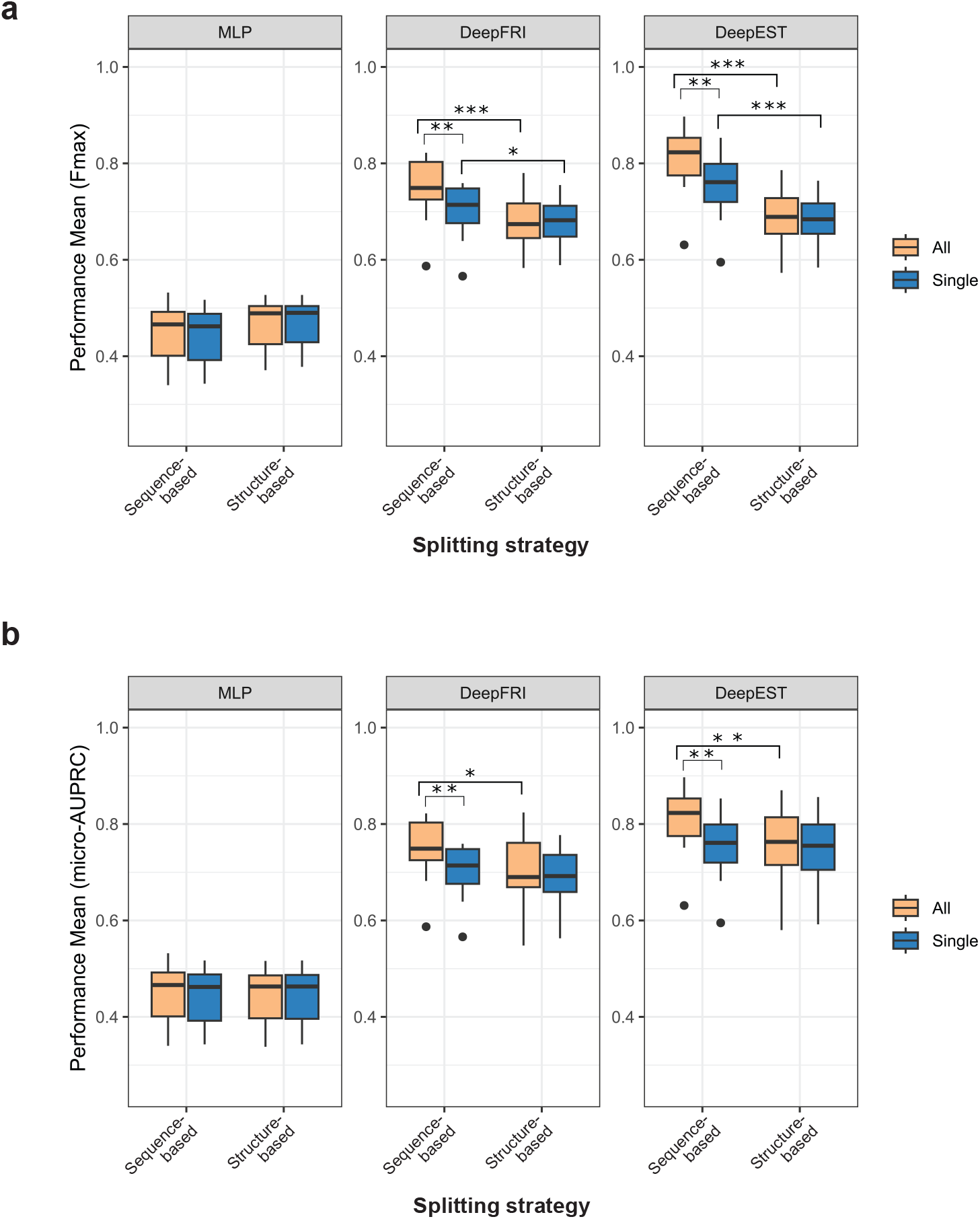
Boxplot comparing sequence (Seq) and structure (Str) accounting performance for three models: MLP, DeepFRI, and DeepEST. Performance is measured as **a** *F*_max_ and **b** *micro*-AUPRC. The models are displayed as separate facets in the specified order (MLP, DeepFRI, DeepEST). Each box represents the distribution of mean values under two conditions (colored bars): “All” (orange) and “Single” (blue). Error bars show the range of performance, and the y-axis is restricted to a range of 0.25 to 1 for clarity. Statistically significant results from the Wilcoxon rank-sum test (unpaired) are indicated by “*” (pvalue *<* 0.05), “**” (pvalue *<* 0.01) and “***” (pvalue *<* 0.001).

**Fig. S.9:**
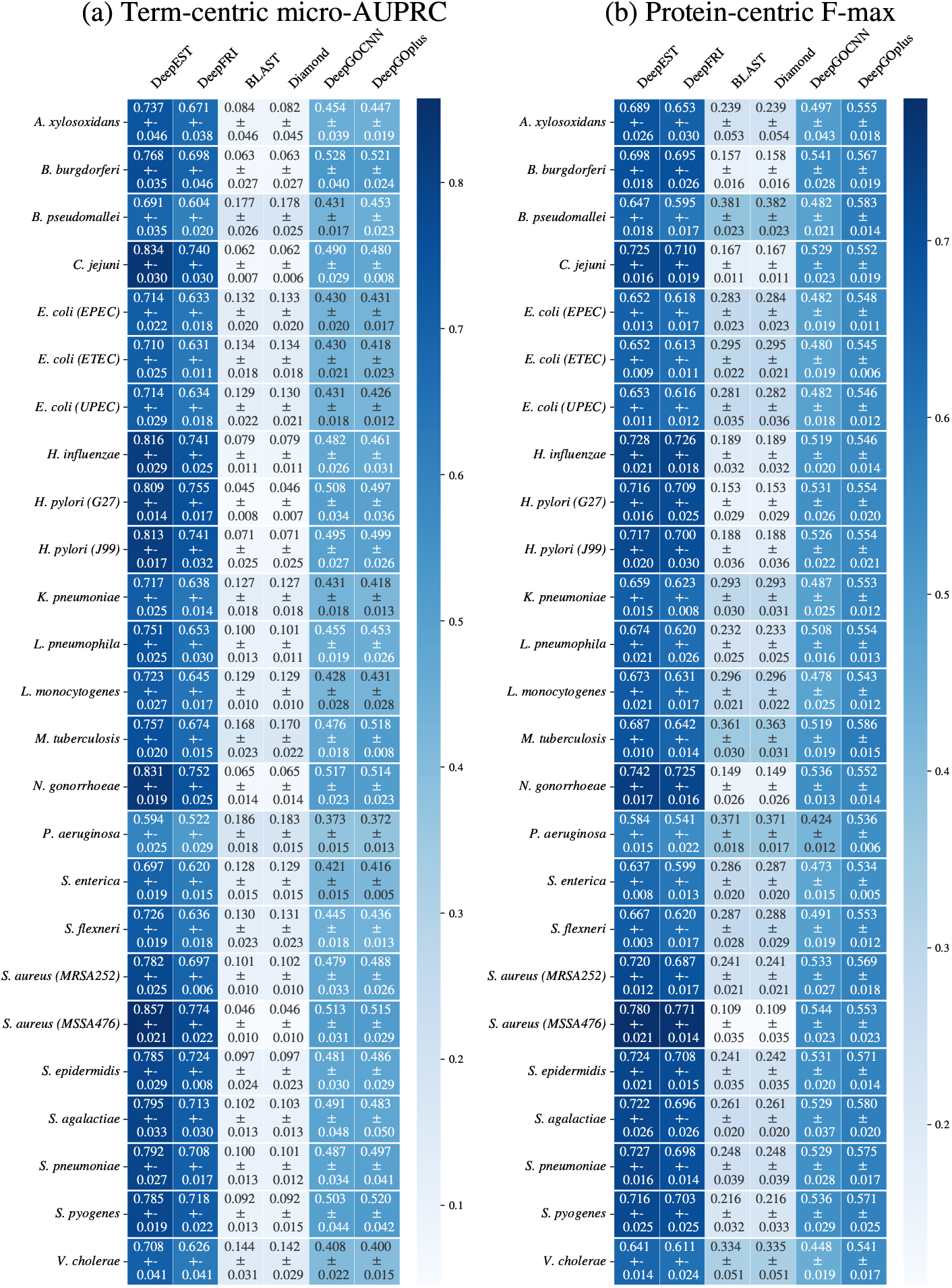
We report the results, as average ± standard deviation across the five test sets, for DeepEST and the comparison partners (DeepFRI, BLAST, Diamond, DeepGOCNN, DeepGOplus): (a) *micro-*AUPRC (b) *F*_max_.

**Fig. S.10:**
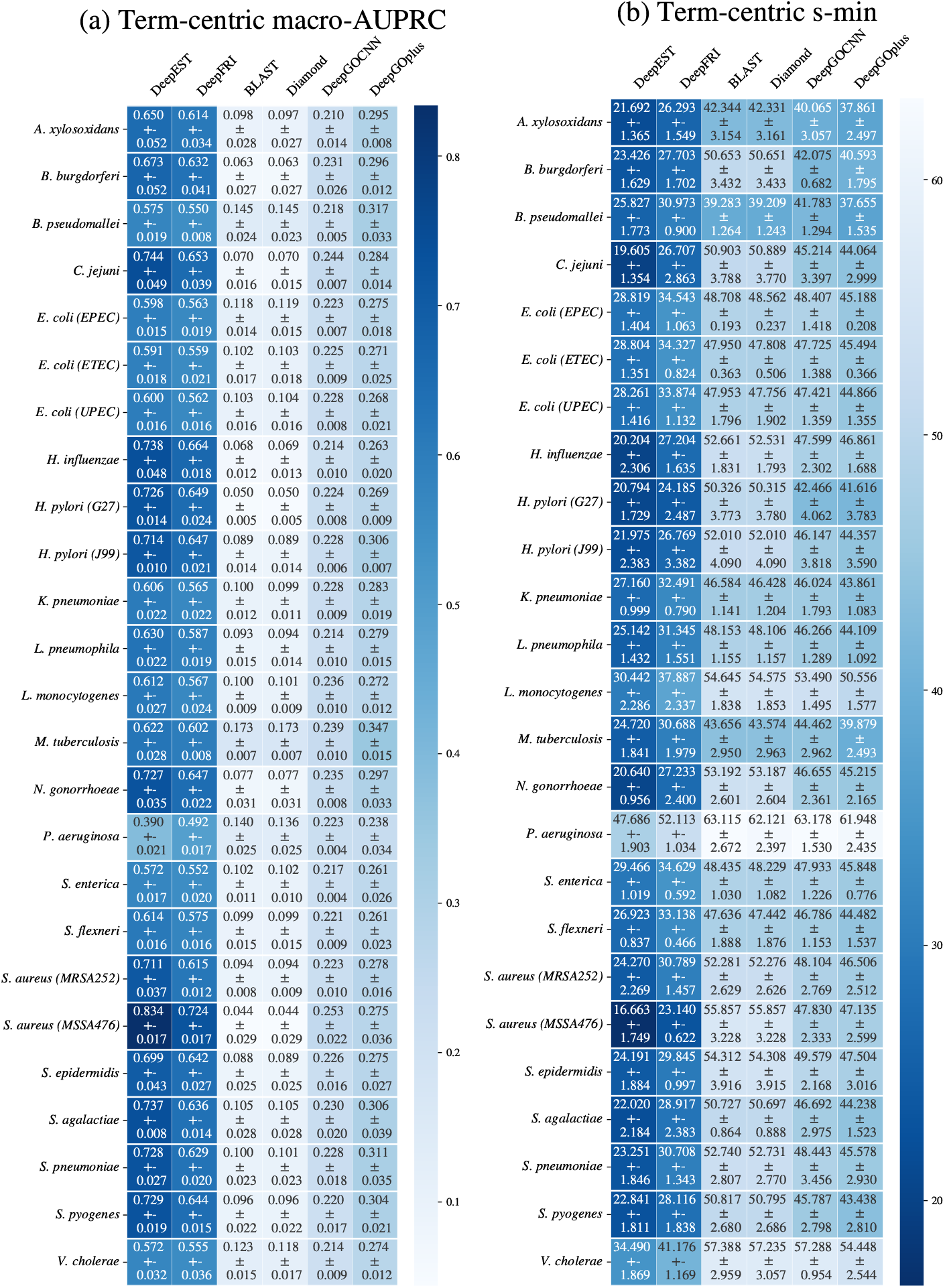
We report the results, as average ± standard deviation across the five test sets, for DeepEST and the comparison partners (DeepFRI, BLAST, Diamond, DeepGOCNN, DeepGOplus): (a)*macro-*AUPRC (b) *s*_min_. Note that for *s*_min_, lower values represent a better performance in comparison to higher values.

**Fig. S.11:**
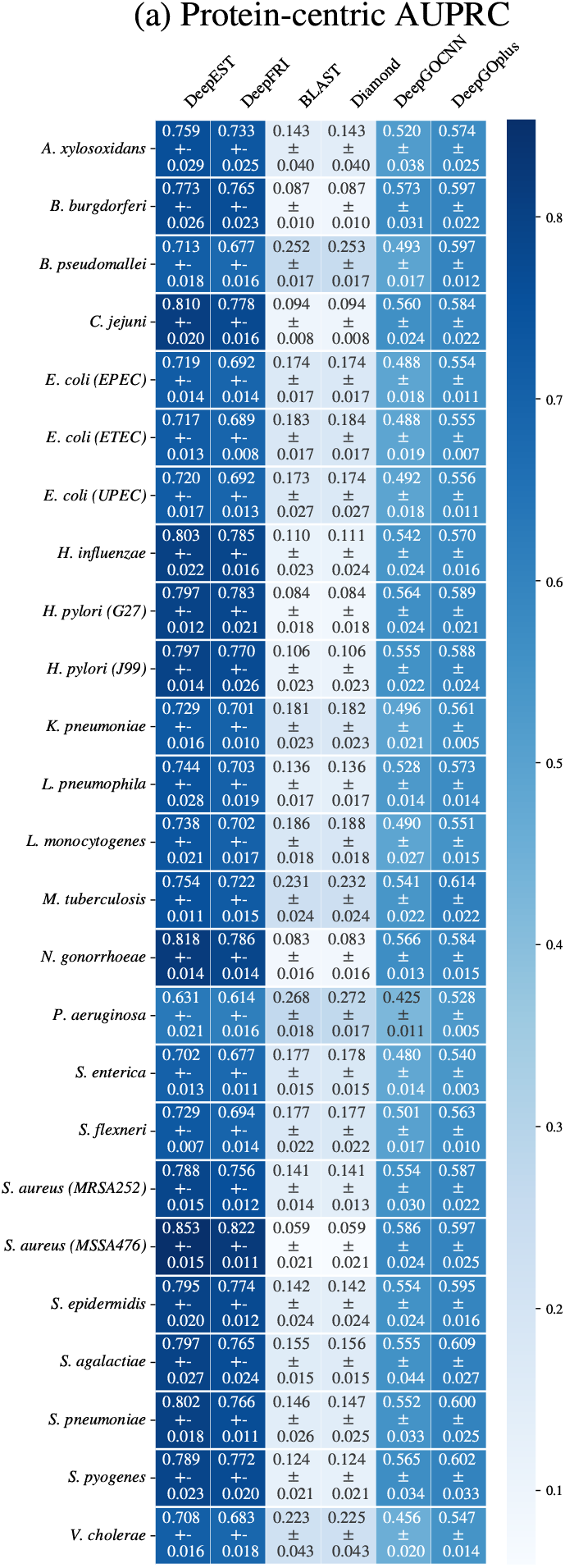
We report the results, as average ± standard deviation across the five test sets, for DeepEST and the comparison partners (DeepFRI, BLAST, Diamond, DeepGOCNN, DeepGOplus): (a) Protein-centric AUPRC.

**Fig. S.12:**
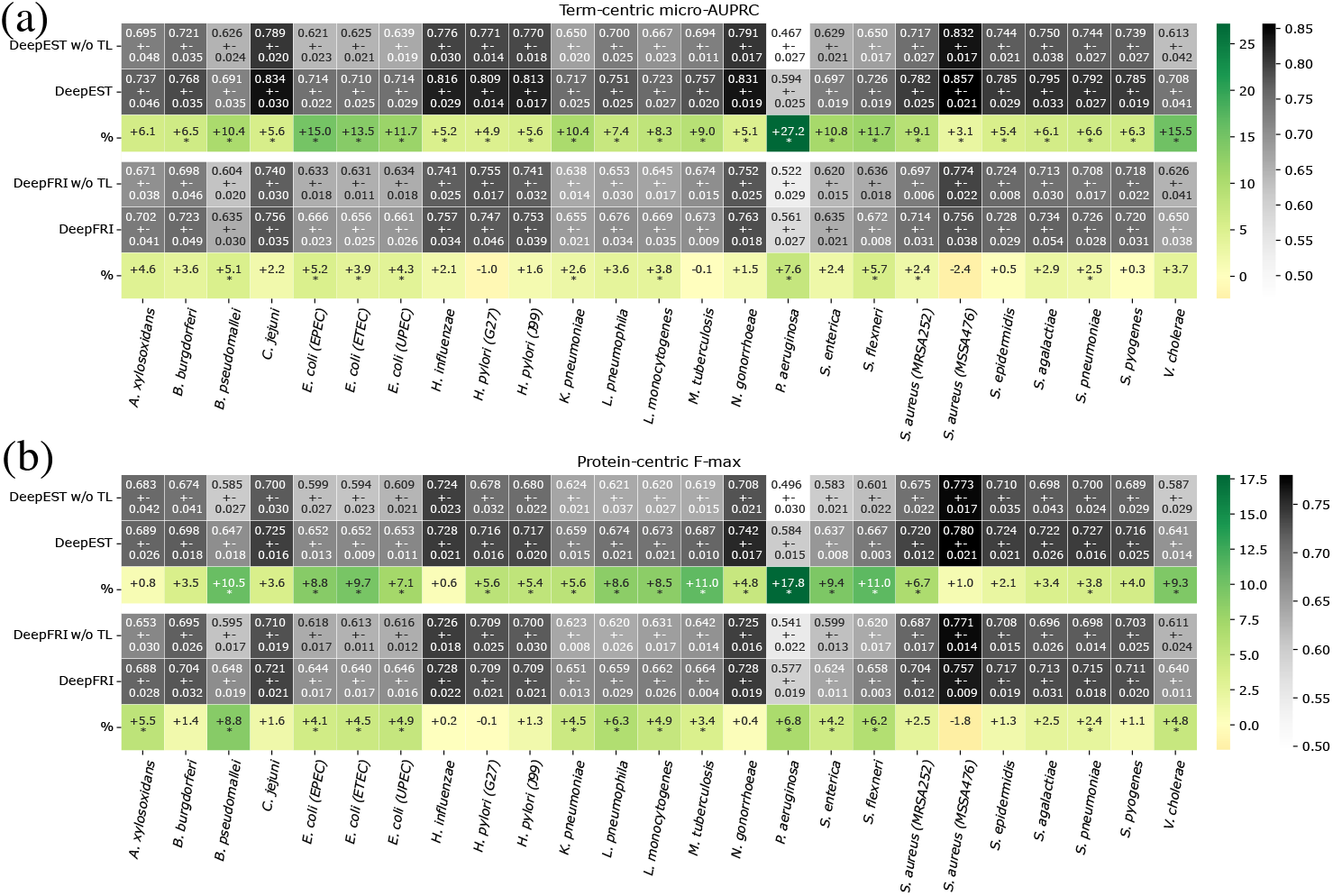
Comparison between not using (w/o TL) and using our transfer learning technique on both DeepEST and DeepFRI. We report the results as average ± standard deviation across the five test sets in terms of (a) *micro-*AUPRC and *F*_max_. We additionally report the percentage change of performance when using transfer learning in comparison to not using it. These percentages are labeled with an “*” when the difference between the two averages exceeds one standard deviation. Each column represents a species.

**Fig. S.13:**
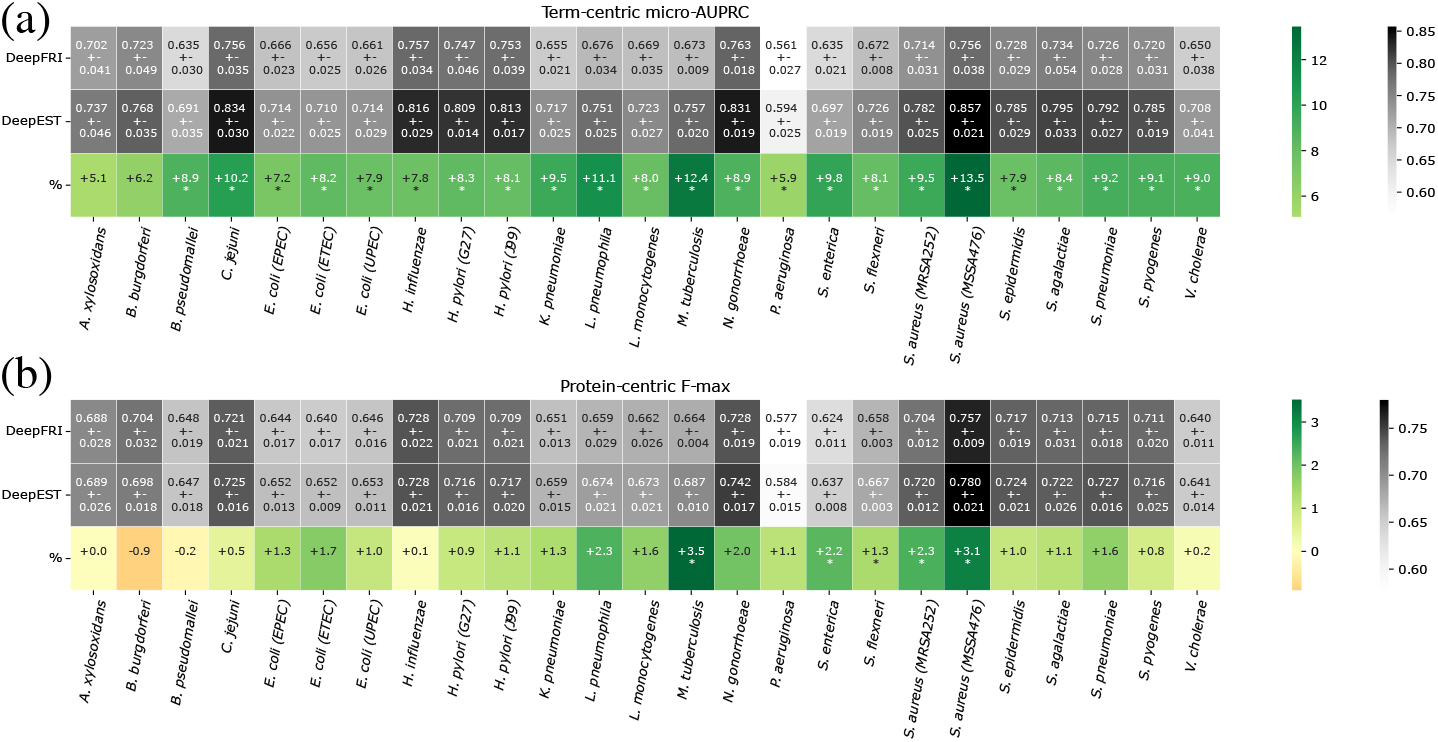
Comparison between *f*_s_ (i.e., DeepFRI with the use of transfer learning) and DeepEST in terms of (a) *micro-*AUPRC and (b) *F*_max_ ™ score. The third line, labeled with “%”, represents the percentage change in performance of DeepEST in comparison to DeepFRI. This percentage is marked with an asterisk when this change exceeds one standard deviation from the average performance of DeepFRI. These are the results on the test sets.

**Fig. S.14:**
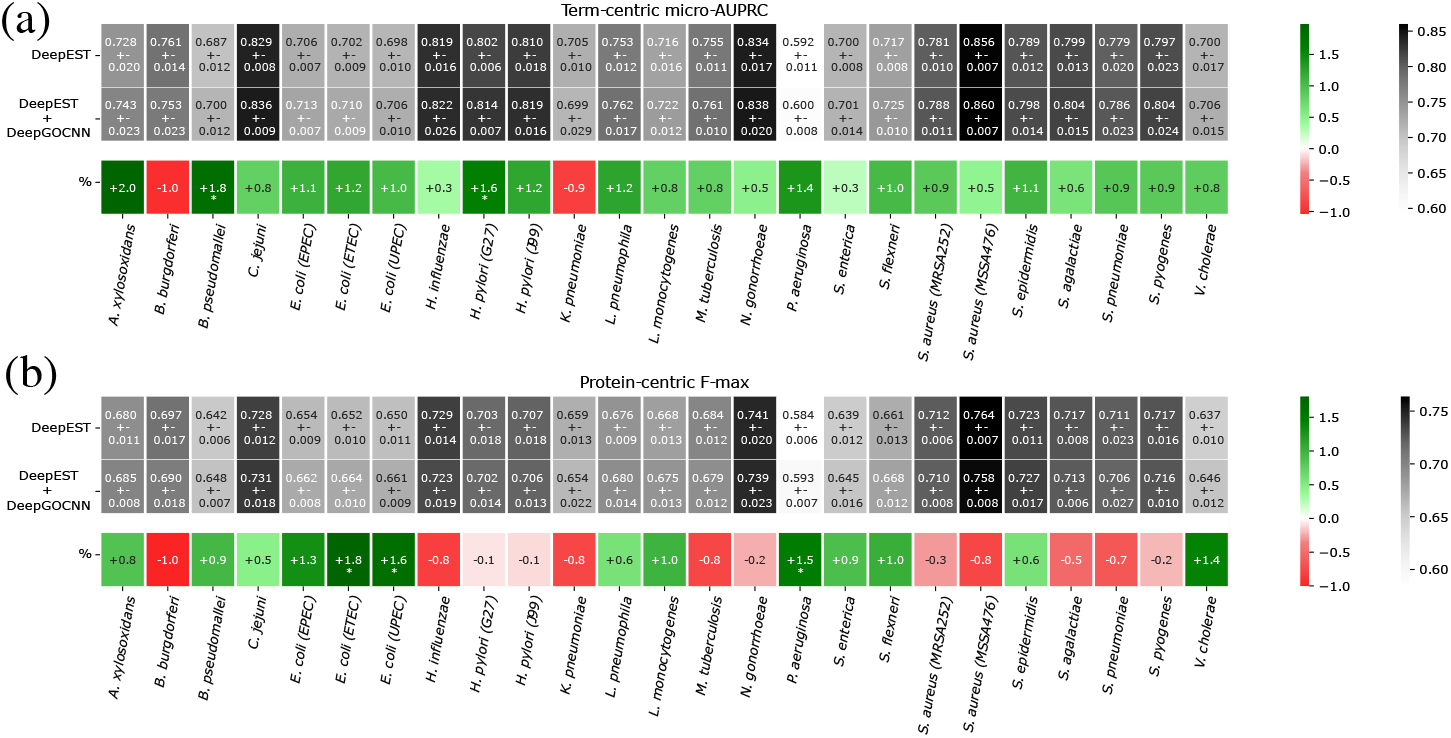
Comparison between DeepEST and DeepEST combined with DeepGOCNN (to include the linear sequence information) in terms of average (a) *micro*-AUPRC and (b) *F*_max_ on the five *validation* sets, with *structure*-based splits. % represents the percentage change in performance when using DeepEST + DeepGOCNN in comparison to DeepEST, with an “*” when the difference between the two averages exceeds one standard deviation. Each column represents a species.

**Fig. S.15:**
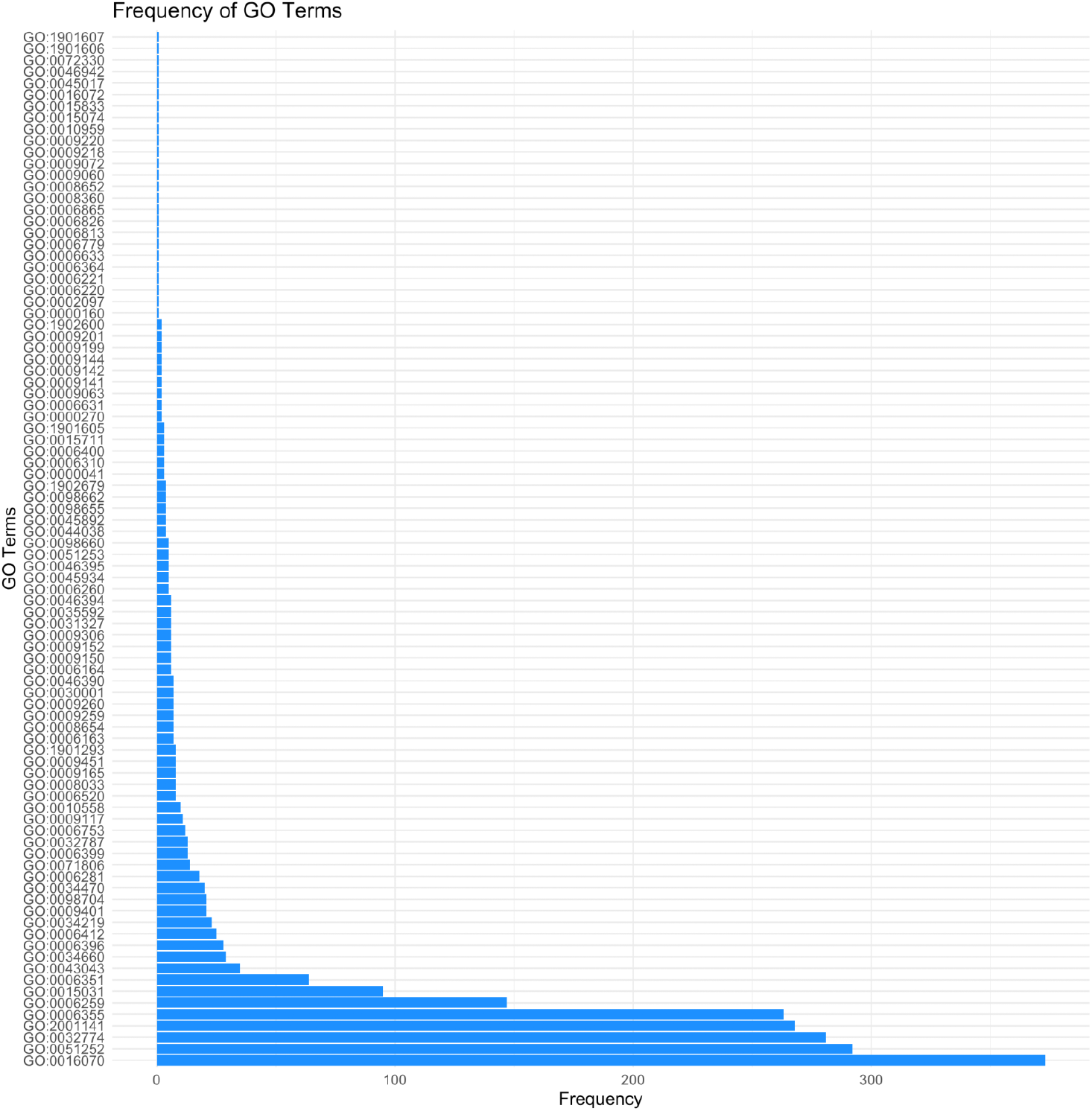
Histogram showcasing connections between GO terms and their corresponding target genes at depths greater than 5.

**Fig. S.16:**
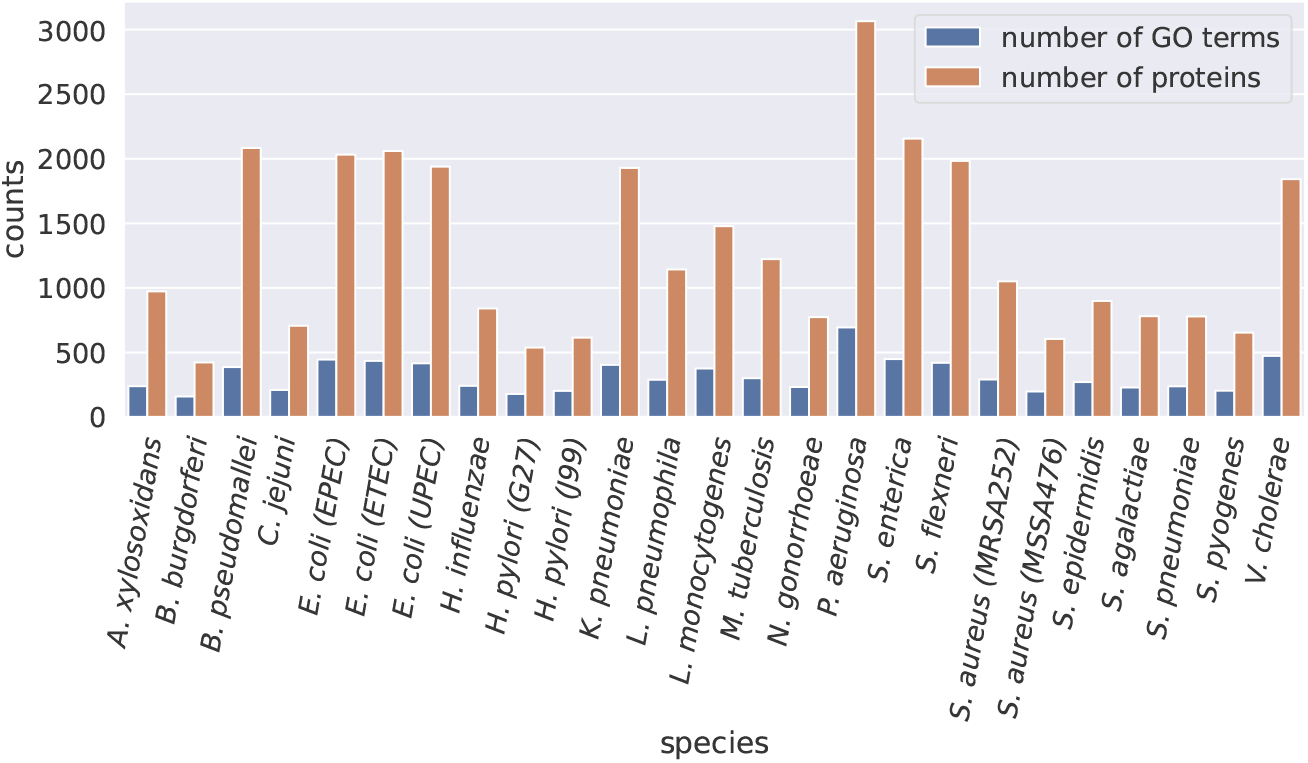
Number of annotated proteins and number of GO terms for each species obtained after having filtered out: (i) GO terms annotated for less than 10 proteins and (ii) proteins presenting no annotations.

**Fig. S.17:**
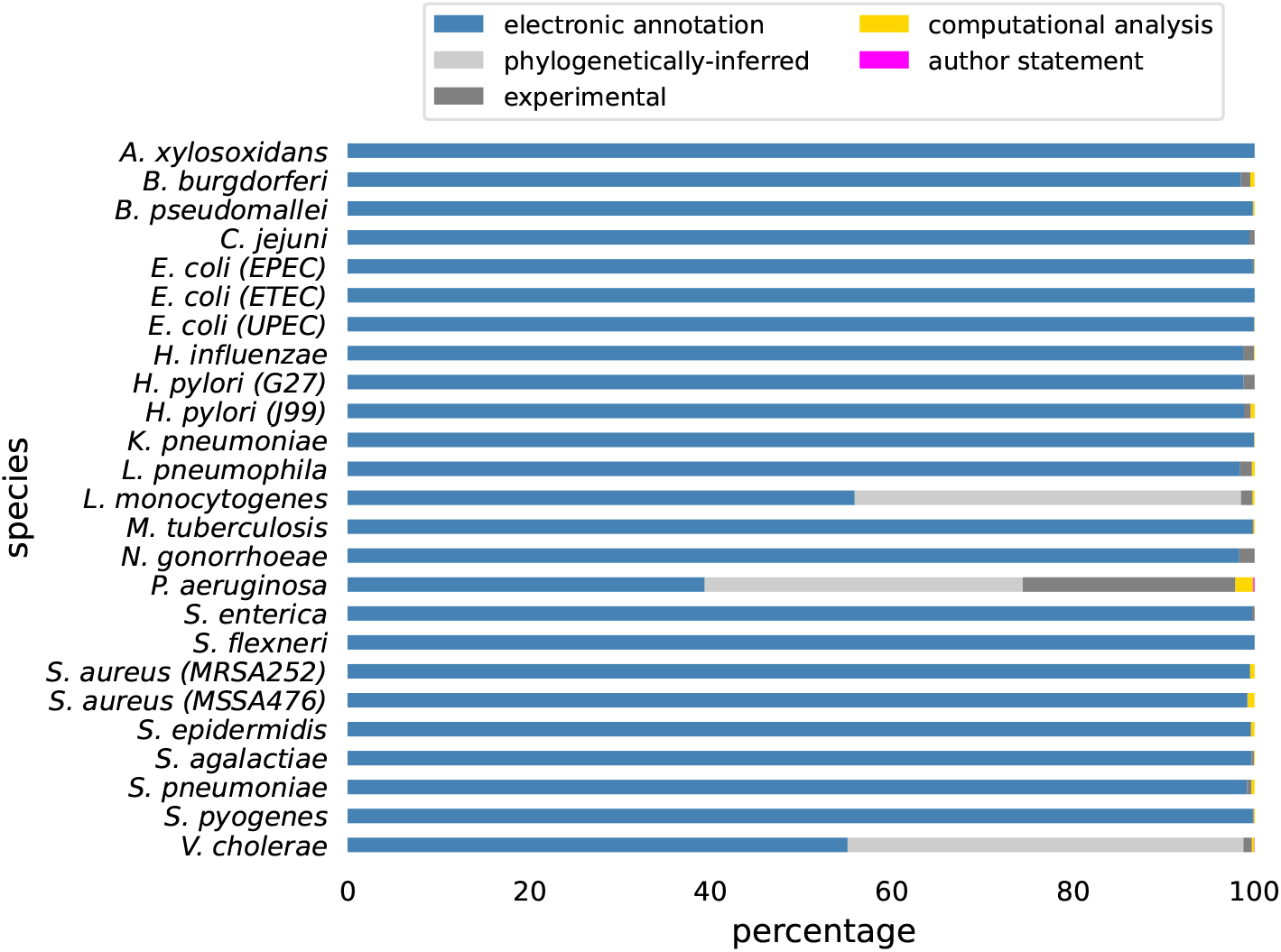
Distribution of the GO terms annotation categories for the selection of 25 species. Each annotation category maps to specific evidence codes, namely: (i) electronic annotation to IEA, (ii) phylogenetically inferred to IBA, (iii) experimental to EXP, IDA, IEP, IGI, IMP and IPI, (iv) computational analysis to IGC and ISS, and (v) author statement to NAS and TAS.

**Fig. S.18:**
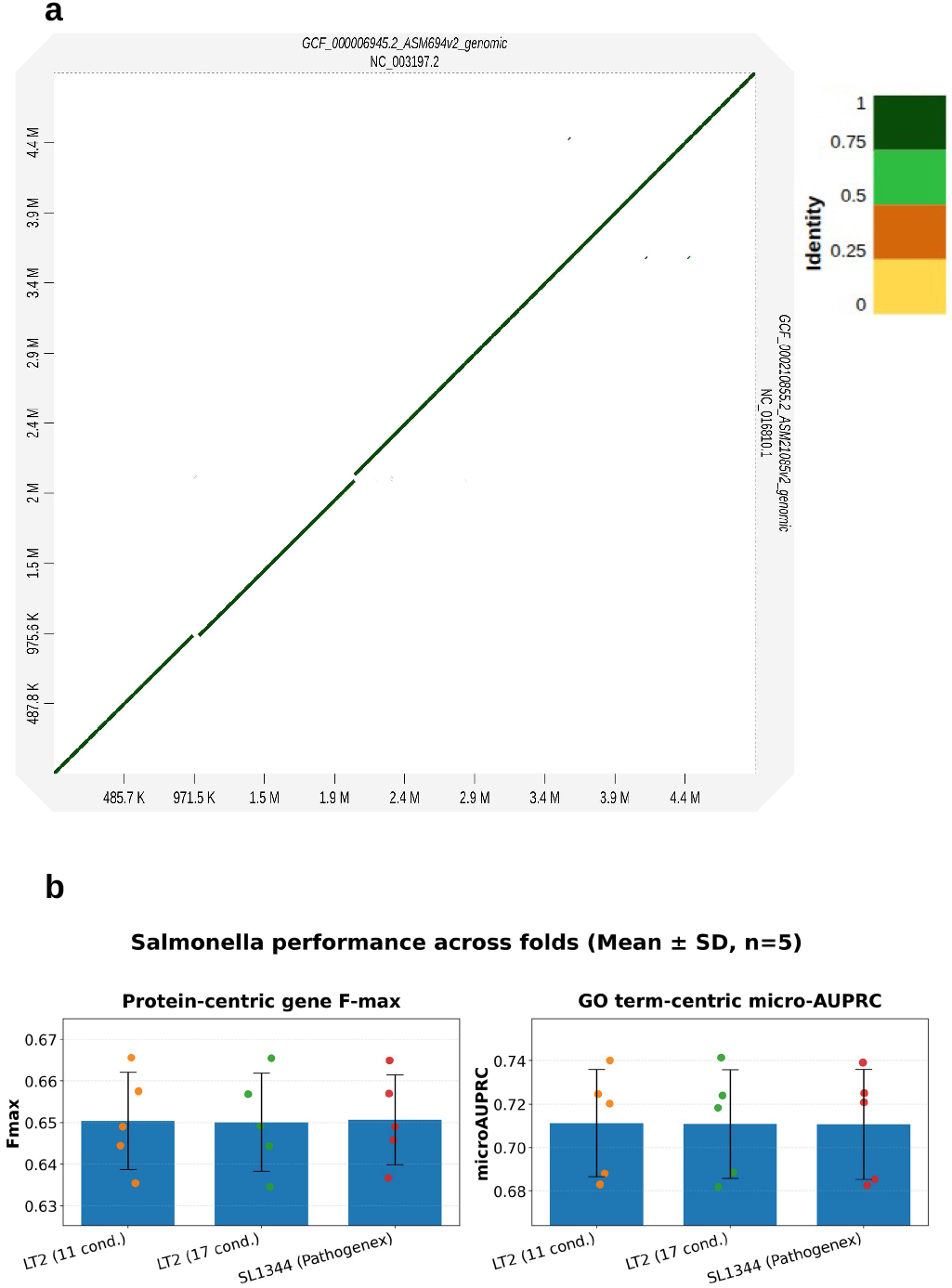
DeepEST performance using user-defined transcriptomic datasets in *Salmonella enterica* LT2. (a) Whole-genome alignment dot plot comparing *Salmonella enterica* serovar Typhimurium SL1344 (NC 016810.1) and LT2 (NC 003197.2). The strong diagonal pattern indicates high nucleotide identity and near-complete conservation of genome organization with minimal rearrangements, supporting transferability of genome-location features between the two strains. Color scale represents sequence identity. (b) DeepEST performance comparison across five cross-validation folds (mean ± SD, n = 5) for SL1344 using PATHOgenex transcriptomic data and LT2 using independently derived transcriptomic profiles. LT2 expression features were computed from the iModulon/Salmonella compendium using Δ*z*-scores relative to exponential growth controls. Results are shown for protein-centric F-max (left) and GO term–centric micro-AUPRC (right). Comparable performance across SL1344 and LT2 demonstrates that DeepEST can incorporate user-defined transcriptomic datasets without loss of predictive accuracy.

